# Spatially resolved transcriptomic identification of thousands of neurons recorded *in vivo*

**DOI:** 10.64898/2026.05.15.725413

**Authors:** Isabelle Prankerd, Maxwell Shinn, Paul C. Shuker, Zhiyao Zhou, Reilly Tilbury, Joshua A. M. Duffield, Christina A. Maat, Dimitris Nicoloutsopoulos, Anne Ritoux, Paula V. Maglio Cauhy, David Orme, Mathieu Bourdenx, Karen E. Duff, Stéphane Bugeon, Yoh Isogai, Kenneth Harris

## Abstract

Transcriptomics has transformed our understanding of the brain, but assigning transcriptomic identities to neurons recorded in vivo remains challenging at scale. Existing platforms can pair transcriptomic identity with two-photon calcium imaging in small populations of approximately 100 neurons, but they require recorded cells to be sparse and therefore cannot be applied to large population recordings. Here, we present coppaFISH 3D, a spatially resolved transcriptomics method, and CASTalign, an in silico alignment framework, which together enable transcriptomic identification of thousands of simultaneously recorded cells. coppaFISH 3D detects hundreds of genes in thick 50μm fixed sections while preserving tissue integrity, enabling both 3D registration to in vivo imaging and integration with immunofluorescence labelling. The platform is fully powered by open chemistry and open source software, runs on commodity hardware, and can be performed at very low cost per section. It therefore enables transcriptomic identification of recorded neurons at scale, making it possible to study how transcriptomic identity shapes activity in neural populations.

## Introduction

A neuron’s gene expression profile contains rich information about its morphology, connectivity, and physiology^1–7^. Yet, relating gene expression to a neuron’s role in neural circuits remains challenging: gene expression is typically measured *ex vivo*, whereas neuronal activity must be measured in living animals. One promising approach is to record neuronal activity *in vivo* using two-photon calcium imaging, measure gene expression in the imaged tissue *ex vivo*, and then register the two measurements together using image alignment^8–19^. Imaging-based spatially resolved transcriptomic methods, which can measure expression of hundreds of genes simultaneously in tissue sections^20–34^, provide a particularly useful way to do this.

Despite several successes, this strategy remains difficult to scale beyond approximately 100 recorded neurons (Supplementary Table 1). Insufficient cell counts are especially limiting in brain regions with highly diverse cell populations, where only a few neurons of each transcriptomic type may be recorded in any session. They also make it difficult to understand cell type specificity in population codes, as these analyses require large numbers of neurons per transcriptomic type.

Registration of *in vivo* images to *ex vivo* tissue sections is the central obstacle in scaling the method. *Ex vivo* brain processing induces nonlinear tissue warping, and even small distortions can defeat automated alignment methods. The problem is exacerbated because most existing methods use unfixed tissue cut into sections that are around the size of a single neuron, both of which increase tissue warping.

Previous methods have therefore typically relied on hand-matching small numbers of sparsely-labelled neurons, one by one^8,10–18,35^. This is slow, and in dense fields of view, may be insufficient to unambiguously identify neurons. New experimental and computational methods are needed to enable more efficient registration.

Here, we introduce coppaFISH 3D, a spatially-resolved transcriptomics platform, and CASTalign, a registration method to link gene expression patterns to *in vivo* neuronal recordings. coppaFISH 3D can identify hundreds of genes simultaneously in 3D fixed tissue sections while preserving tissue integrity for post hoc immunofluorescence (IF) antibody staining. CASTalign uses this 3D structure to semi-automatically register thousands of neurons between *in vivo* calcium imaging and *ex vivo* transcriptomics, requiring only minimal changes to existing experimental protocols and less than one hour per section of human involvement. Together, they improve mammalian *in vivo* cell type identification by 1-2 orders of magnitude (Supplementary Table 1) while being fully open source, with a running cost of only £250 per section after a one-time equipment purchase of approximately £320,000. coppaFISH 3D and CASTalign make high throughput studies of gene expression, cell type, and neuronal activity possible and accessible.

Our platform links molecular cell identities to *in vivo* neuronal activity or post hoc protein labelling (Figure 1a) through a workflow that spans from *in vivo* imaging to spatially-resolved transcriptomics (Figure 1b). We first present coppaFISH 3D as a 3D spatially resolved transcriptomics protocol for thick PFA-fixed sections, together with the gene calling and cell calling analyses that convert sequential volumetric images into molecular cell identities. We then validate transcript detection and cell type assignment, and show that the same processed tissue can be linked to post hoc IF labelling. Finally, we present CASTalign, a software tool to register coppaFISH 3D to *in vivo* two-photon calcium recordings, showing that thousands of recorded neurons can be paired with molecular identities. Throughout, we emphasise the features needed for experiments on precious *in vivo* imaged tissue: low warping, recoverability after technical faults and preservation of tissue integrity. All software and protocols are freely available and extensively documented, and the platform is modular so that individual components can be iteratively improved or repurposed for other work-flows.

**Figure 1:**
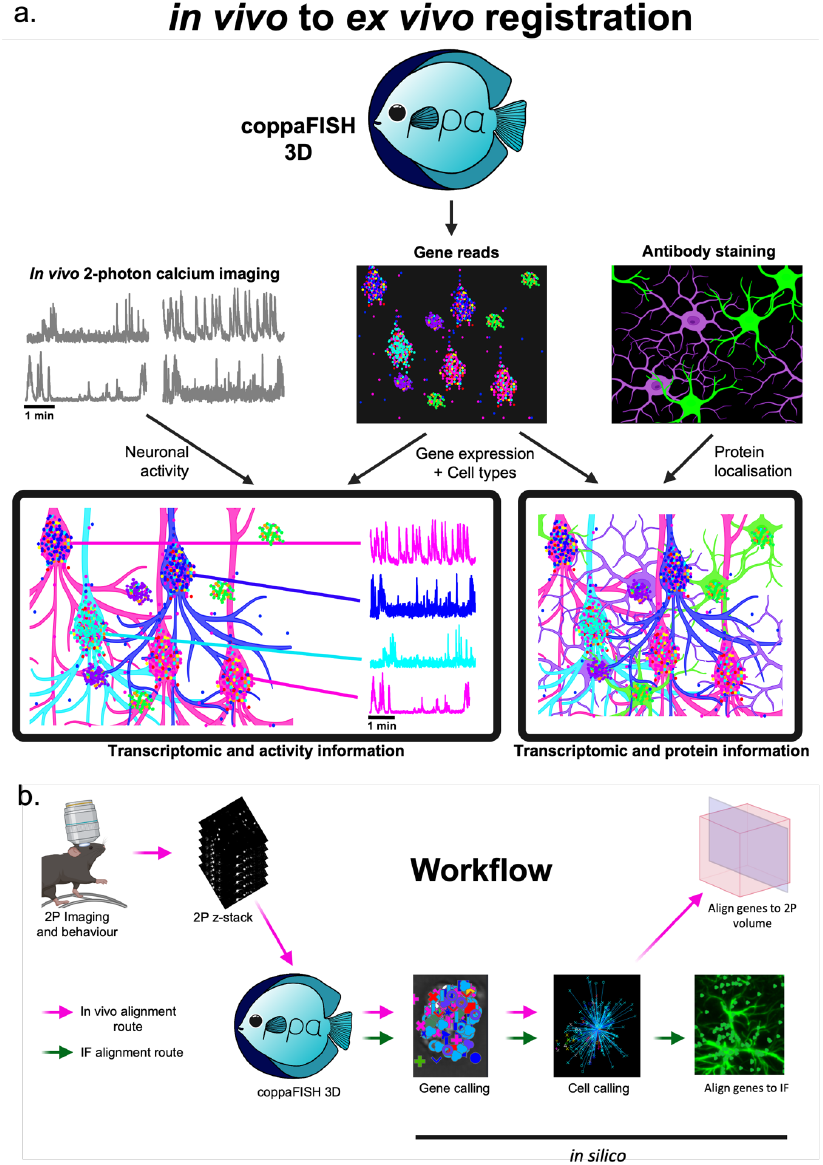
coppaFISH 3D and CASTalign overview. (a) coppaFISH 3D detects genes *in situ* which enables identification of transcriptomic subtypes. This can be combined with *in vivo* two-photon calcium imaging to link cell types and gene expression to neuronal activity or combined with antibody staining to link them to protein expression. (b) Schematic of the full workflow: mice undergo two-photon imaging and acquisition of a reference z-stack, followed by perfusion, sectioning, coppaFISH 3D, gene calling, cell typing, and alignment either back to the *in vivo* imaging volume or to post hoc antibody staining. Figure created with BioRender.com.

## Results

### Overview

Our platform links molecular cell identities to *in vivo* neuronal activity or post hoc protein labelling (Figure 1a) through a workflow that spans from *in vivo* imaging to spatially-resolved transcriptomics (Figure 1b). We first present coppaFISH 3D as a 3D spatially resolved transcriptomics protocol for thick PFA-fixed sections, together with the gene calling and cell calling analyses that convert sequential volumetric images into molecular cell identities. We then validate transcript detection and cell type assignment, and show that the same processed tissue can be linked to post hoc IF labelling. Finally, we present CASTalign, a software tool to register coppaFISH 3D to *in vivo* two-photon calcium recordings, showing that thousands of recorded neurons can be paired with molecular identities. Throughout, we emphasise the features needed for experiments on precious *in vivo* imaged tissue: low warping, recoverability after technical faults and preservation of tissue integrity. All software and protocols are freely available and extensively documented, and the platform is modular so that individual components can be iteratively improved or repurposed for other workflows.

### coppaFISH 3D

coppaFISH 3D is an imaging-based spatially resolved transcriptomics method optimised for 50µm thick, PFA-fixed tissue sections. This format reduces tissue warping, increases throughput, and facilitates alignment with *in vivo* neuronal recordings. If technical faults occur, tissue can be reimaged, providing robustness when working with precious brain tissue. Furthermore, coppaFISH 3D is customisable and affordable: the necessary hardware cost approximately £320k at the time of purchase, and the cost per section is about £250 for a panel of 124 genes and £265 for a panel of 320 genes (Supplementary table 2 and 3).

The coppaFISH 3D workflow has six main steps: RNase-free perfusion and sectioning, permeabilisation, transcript amplification, optional fluorescent protein imaging for *in vivo* alignment, and sequential imaging for gene identification (Figure 2a).

**Figure 2.**
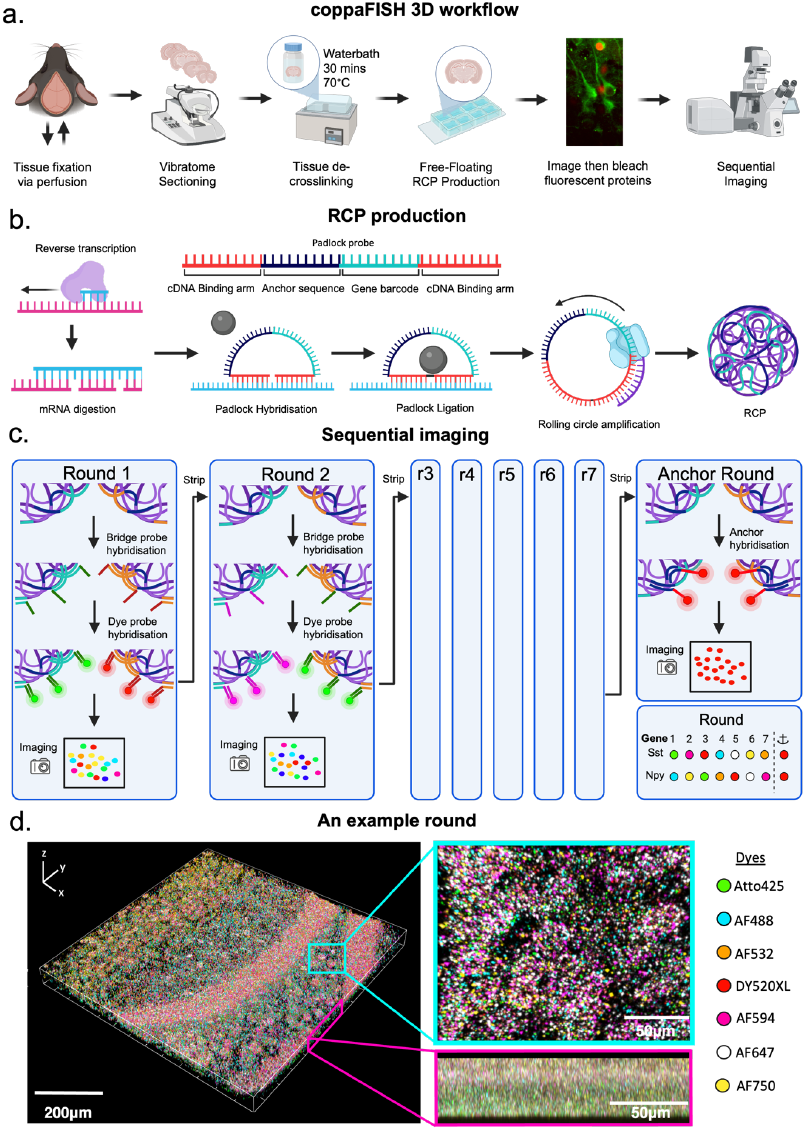
coppaFISH 3D workflow for transcript detection in 50 μm fixed tissue sections. (a) Experimental workflow, from transcardial perfusion and vibratome sectioning through tissue de-crosslinking, free-floating rolling-circle-product (RCP) generation, fluorescent protein imaging and bleaching, and sequential confocal imaging. (b) Schematic of RCP production. Individual mRNA molecules of genes of interest are reverse transcribed. Padlock probes containing two gene binding segments, an anchor sequence segment, and a barcode segment are hybridised to genes and amplified to form RCPs. These RCPs spatially tag individual mRNA molecules and make them visible as spots under a confocal microscope. (c) Sequential imaging strategy. In each sequencing round, bridge and dye probes assign a colour to each gene via a unique barcode; after seven rounds, an anchor round labels all RCPs to provide a common reference for registration and decoding. These rounds collectively create a combinatorial code capable of identifying hundreds of genes. (d) Example imaging data from one round, showing a 3D tissue volume and higher-magnification views in orthogonal planes, together with the dye set used for combinatorial readout. Figure a-c created with BioRender.com.

Transcardial perfusion and vibratome sectioning prepare tissue while preserving mRNA integrity. To avoid RNA degradation, perfusion is performed using RNase-free reagents, tools, fixatives, and buffers. Sections are cut at 50µm and can be any orientation, but we recommend tangential to the *in vivo* imaging plane when available.

Permeabilisation exposes mRNA to reagents and decrosslinks the fixed tissue. To enable registration to *in vivo* imaging, we use a protocol that preserves fluorescent proteins: water bath treatment in a citrate buffer for 30 minutes at 70°C. When registration is not required, sections are treated at 80°C to bleach fluorescent proteins. This water bath method was selected from several alternatives as we found it to best increase transcript accessibility (Supplementary Figure 1). After permeabilisation, sections are dried and stored in well plates at −80°C; RNA integrity was preserved in even our oldest samples which are 3 years post sectioning.

Transcript amplification produces the rolling-circle products (RCPs) used to visualise individual transcripts by light microscopy. We modified the coppaFISH^18^ chemistry for thick free-floating sections in well plates rather than coverslip-mounted tissue, allowing reagents to enter from both sides and improving access across the section depth. In brief, gene-specific primers first hybridise to target mRNAs and are extended by reverse transcription to generate cDNA. After the RNA template is digested, padlock probes^36^ hybridise to the cDNA and are ligated. Each padlock probe contains two cDNA-binding arms, a gene-specific barcode, and a common anchor sequence. Rolling-circle amplification of the ligated padlock then generates an RCP at the transcript location (Figure 2b). After this step, the fixed sections are stable enough to move with a paintbrush without damage.

Fluorescent protein imaging identifies fluorescently labelled cells for alignment with *in vivo* imaging, without interfering with later *ex vivo* sequential imaging for the transcriptomic protocol. Sections are stained with DAPI, imaged in 3D for fluorescent proteins (e.g. GCaMP and dTomato), and then chemically bleached with a solvent and strong detergent so that the same tissue can undergo sequential imaging without interference from the fluorescent proteins.

Sequential imaging then converts the RCPs into a 3D combinatorial readout for gene identification. Sections undergo eight rounds of automated confocal imaging coupled to automated fluidics: seven sequencing rounds followed by one anchor round (Figure 2c). In each sequencing round, round-specific bridge probes bind the gene-specific barcode sequence in each RCP and recruit one of seven dye probes, assigning that gene a colour for that round according to the combinatorial code. After volumetric imaging, the bridge and dye probes are stripped, and the next round is performed. Across the first seven rounds, the sequence of colours observed at each RCP identifies the corresponding gene. The eighth round is not part of the combinatorial code; instead, an anchor probe labels all RCPs through the shared anchor sequence, providing a common reference for registration across rounds and for decoding. An example imaging round is shown in Figure 2d, and the four-camera imaging setup used to increase acquisition speed is shown in Supplementary Figure 2.

### Gene Calling

Gene calling is a data processing step that converts the eight sequential imaging rounds into transcript identities. The full gene-calling pipeline (Figure 3c; Supplementary Figure 3) begins by correcting volumetric images for vignetting and the microscope point-spread function and then detects bright, unambiguous RCPs. It uses these detections together with DAPI and anchor images to register sequencing rounds and colour channels into the anchor coordinate system with sub-pixel accuracy, correcting for drift, tissue deformation and chromatic aberration. Adjacent fields of view are then stitched into a single tissue volume. Gene identity is decoded from the registered fluorescence vector at each location, with learned corrections for spectral overlap, brightness differences, and round-, tile- and channel-specific intensity variation. Finally, orthogonal matching pursuit (OMP) separates spatially overlapping RCPs in the full 3D voxel data, yielding transcript identities, positions and confidence scores for downstream cell calling. A full description of the algorithm, implemented as a dedicated, modular Python package, can be found in the Methods.

**Figure 3.**
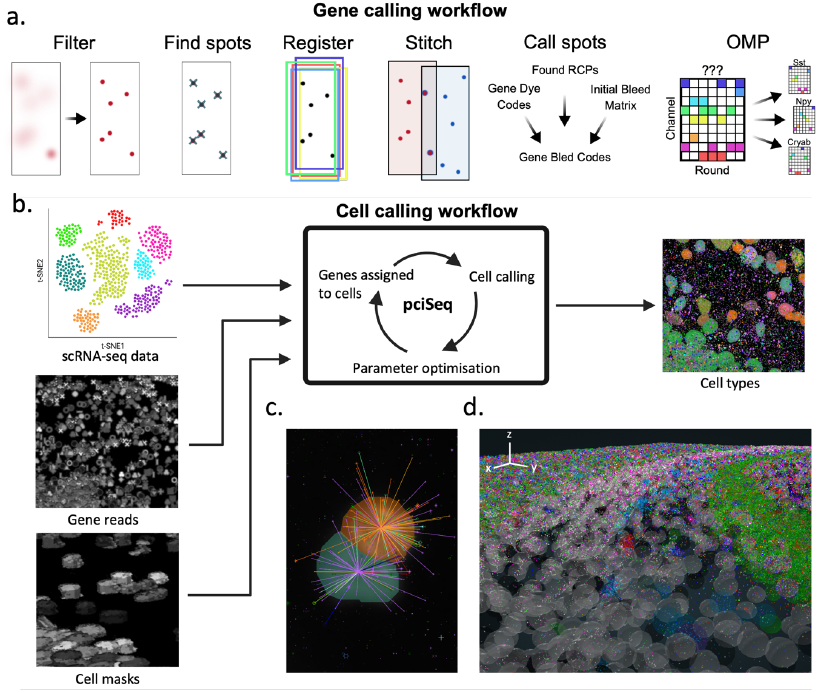
Analysis pipeline for gene calling and cell typing in coppaFISH 3D data. (a) Overview of the gene calling pipeline, including image filtering, spot detection, cross-round registration, tile stitching, initial spot calling, and orthogonal matching pursuit (OMP) decoding. (b) Overview of the cell calling workflow, which combines scRNA-seq reference data, detected gene reads, and cell masks to iteratively assign transcripts to cells and infer cell identities. (c) Illustration of probabilistic gene-to-cell assignment based on spatial proximity and expression profiles. (d) Example 3D view showing segmented cells and their inferred transcriptomic identities in tissue. Figure created with BioRender.com.

### Cell Calling

To assign molecular identities to cells, we adapted a probabilistic cell calling algorithm to use the 3D positions of detected genes. Cells are segmented from DAPI or fluorescent protein images (if available) using a custom 3D extension of Cellpose37 (see Methods). We then use variational Bayes inference to iteratively optimise gene-to-cell assignments and cell identities, extending a 2D probabilistic cell calling algorithm38 for 3D tissue sections (Figure 3b; see Methods). Cell identities are derived from existing scRNA-seq taxonomies39,40 and likelihoods incorporate gene identity, spatial distance, and overall gene expression levels in coppaFISH 3D (Figure 3c). The resulting 3D cell-typed volumes can be explored in our online viewer (Figure 3d, https://acycliq.github.io/coppafish3D_hippocampus/).

### Validation of coppaFISH 3D

We validated coppaFISH 3D using four questions: whether transcript counts were reproducible, whether spatial geneexpression patterns matched independent atlas data, whether measured gene expression recovered transcriptomic cell type structure, and whether assigned excitatory types respected known laminar anatomy.

Transcript counts were highly reproducible within and across mice. In two adjacent coronal sections from two wild type mice, transcript counts were strongly correlated within mouse 1 (Pearson correlation r=0.99), within mouse 2 (r=0.99) and between mice (r=0.97) (Figure 4a).

**Figure 4.**
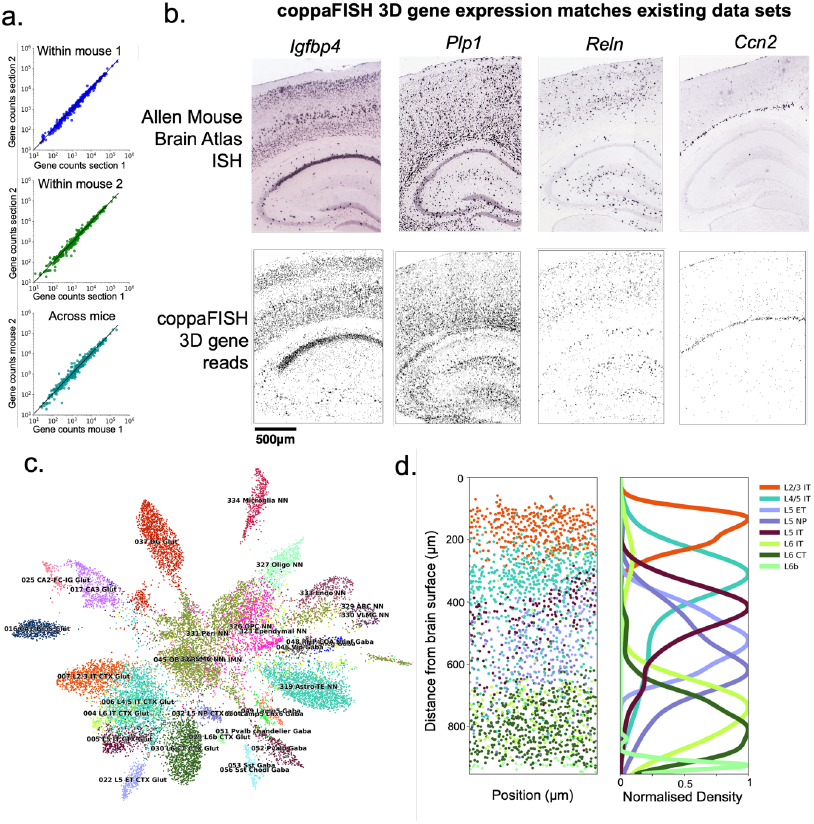
Validation of transcript detection and cell-type assignment in coppaFISH 3D. (a) Gene counts are highly reproducible across adjacent sections from the same mouse and across mice. Scatter plots compare total counts for all detected genes within mouse 1, within mouse 2, and across mice. (b) Spatial expression patterns for representative genes (*Igfbp4, Plp1, Reln*, and *Ccn2*) are consistent between Allen Mouse Brain Atlas^41^ *in situ* hybridisation images (top) and coppaFISH 3D gene reads (bottom). (c) UMAP projection of cells from a section coloured by assigned transcriptomic cell type, showing clustering of related cell classes based on measured expression. (d) Laminar distributions of excitatory cell types in visual cortex, shown as individual cell positions (left) and smoothed density profiles (right), recapitulate expected cortical depth organisation.

The spatial pattern of transcripts was consistent with published data. We compared coppaFISH 3D gene maps from a wild type mouse to the Allen Mouse Brain in situ hybridisation (ISH) atlases41. The spatial patterns were highly similar between the two data sets (Figure 4b; Supplementary Figure 4).

Cell type assignments also recovered expected transcriptomic structure. In a UMAP of all cells from a coppaFISH 3D dataset in a wild type mouse, cells formed clusters within their assigned cell type, with closely related cell types adjacent to one another, despite UMAP having no a priori knowledge of the cell type (Figure 4c). This indicates that transcript and cell type assignment capture the expected transcriptomic structure.

Finally, excitatory neurons in visual cortex showed a spatial organisation consistent with their expected laminar positions, despite the absence of spatial constraints in the gene calling or cell calling algorithms (Figure 4d). This anatomical consistency provides an independent validation of our cell typing pipeline.

### Registration to immunofluorescence antibody staining

coppaFISH 3D is compatible with post hoc IF antibody staining, allowing gene and protein expression to be detected in the same cells. This enables the method to be used for a wider range of applications, such as studying the abnormal protein accumulation in neurodegenerative diseases, as well as *post-hoc* labelling of cells with complex morphologies, such as astrocytes and microglia.

We incorporated IF with coppaFISH 3D by labelling tissue after sequential transcript imaging. Because coppaFISH 3D preserves proteins, processed sections can undergo primary and secondary antibody staining, confocal imaging, and alignment to transcriptomic data using DAPI (Figure 5a; see Methods). Additionally, water bath treatment is a known antigen retrieval method that will typically aid, rather than hinder, antibody compatibility. These features distinguish coppaFISH 3D from platforms that damage proteins or have restrictions on the types of antibodies that may be used.

**Figure 5.**
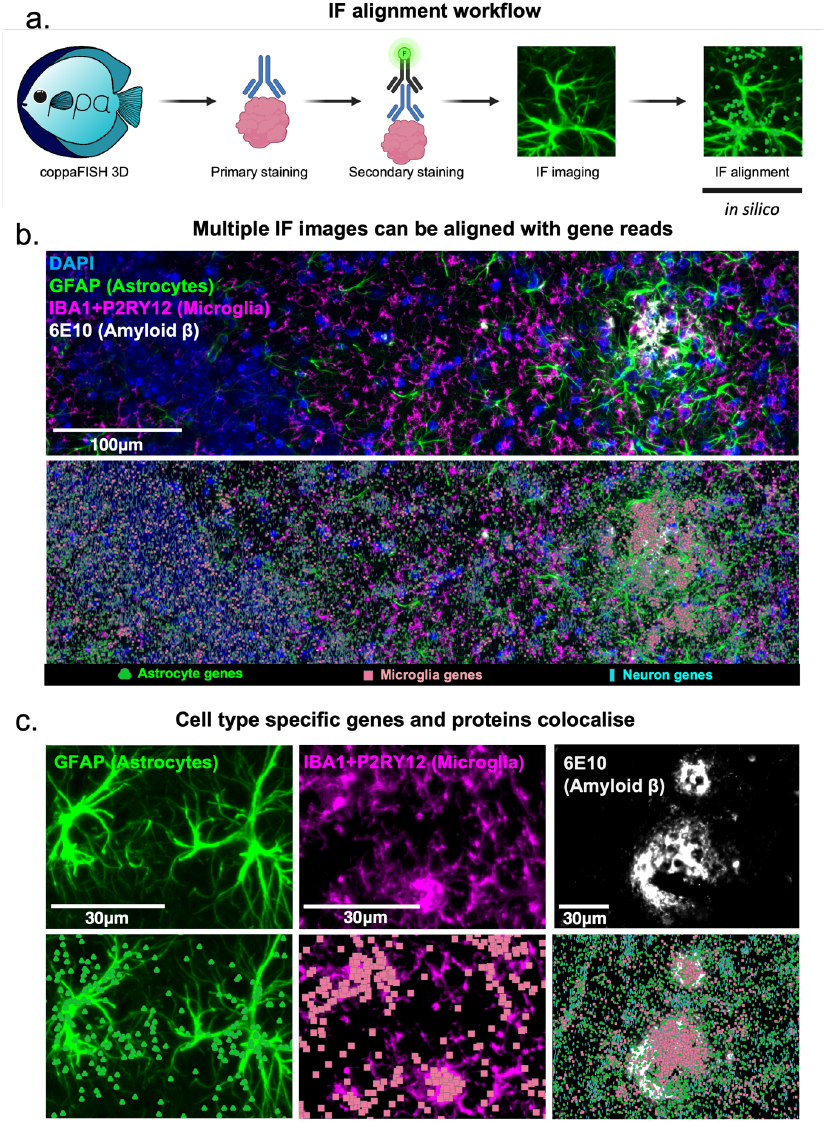
Alignment of coppaFISH 3D data with post hoc IF. (a) Workflow for linking coppaFISH 3D to IF antibody staining after transcript imaging. Following coppaFISH 3D, tissue is immunostained, imaged, and aligned to the transcriptomic data. Figure created with BioRender.com. (b) Example field showing IF staining for astrocytes (GFAP, green), microglia (IBA1/P2RY12, magenta), and amyloid β (6E10, white) with DAPI (blue) in the antibody image (top), and the corresponding aligned transcript reads classified by cell class overlayed (bottom) in an Alzheimer’s disease model mouse. (c) Higher-magnification views illustrating correspondence between antibody signal and transcriptomic readout for astrocytes, microglia, and amyloid-associated cellular organisation.

To validate IF labelling and alignment, we performed coppaFISH 3D followed by IF labelling for several protein markers in a mouse model with Alzheimer-related pathology (MAPT^S305N;Int10+3^ x APP^NL-F^ mouse)^42^. We labelled four proteins: GFAP (an astrocyte marker), IBA1 + P2RY12 (microglial markers) and amyloid β plaques (using the 6E10 antibody). Two rounds of labelling were performed (first round: 6E10, second round: GFAP, IBA1 + P2RY12), and the fluorescent signal was stripped between rounds using the 4i elution protocol^43^. Figure 5b shows the IF rounds aligned to the transcriptomics.

We overlaid transcripts for microglial marker genes (e.g., *Cx3cr1, C1qa, P2ry12* and *Tmem119*) with the IBA1 and P2RY12 antibodies, and astrocyte marker genes (e.g., *Gfap, Slc1a3, Vim* and *Aqp4*) with the GFAP antibody (Figure 5c). Protein and transcripts colocalised for both cell types, validating the alignment and transcript detection procedures.

Finally, amyloid β staining showed microglial transcripts clustered around the plaques, consistent with previous findings that microglia crowd the plaque niche^44,45^.

### Registration to *in vivo* recordings

We developed CASTalign (CoppaFISH, Antibody Staining, and Two-photon alignment) to co-register in vivo multiplane two-photon calcium imaging recordings with ex vivo transcriptomics. CASTalign is a registration framework for aligning hundreds of 3D images into the same coordinate systems. For example, in one recording session in visual cortex, CASTalign paired 3022 recorded neurons to ex vivo segmented neurons and coppaFISH 3D assigned transcriptomic types to 2388 of them. This scale provides access to population-wide activity patterns and large sample sizes for rare cell types.

Registration of 2-photon calcium imaging to transcriptomics is based on fluorescent proteins that are imaged both in vivo and ex vivo. Neurons in the region of interest broadly express the calcium indicator GCaMP, plus a fiducial marker such as dTomato is expressed in sparse cells (such as inhibitory neurons). Calcium imaging can be performed over multiple days and sessions (Figure 6a), and CASTalign will match neurons from all recordings together. In addition to these functional imaging sessions, we acquire a medium-resolution “daily z-stack” on each recording day, and on the first day a high-resolution “reference z-stack” These z-stacks provide a bridge between the functional imaging planes and the ex vivo transcriptomics, allowing imaging planes to be registered to the transcriptomics even when some cells have GCaMP expression too weak to appear in the z-stacks.

**Figure 6.**
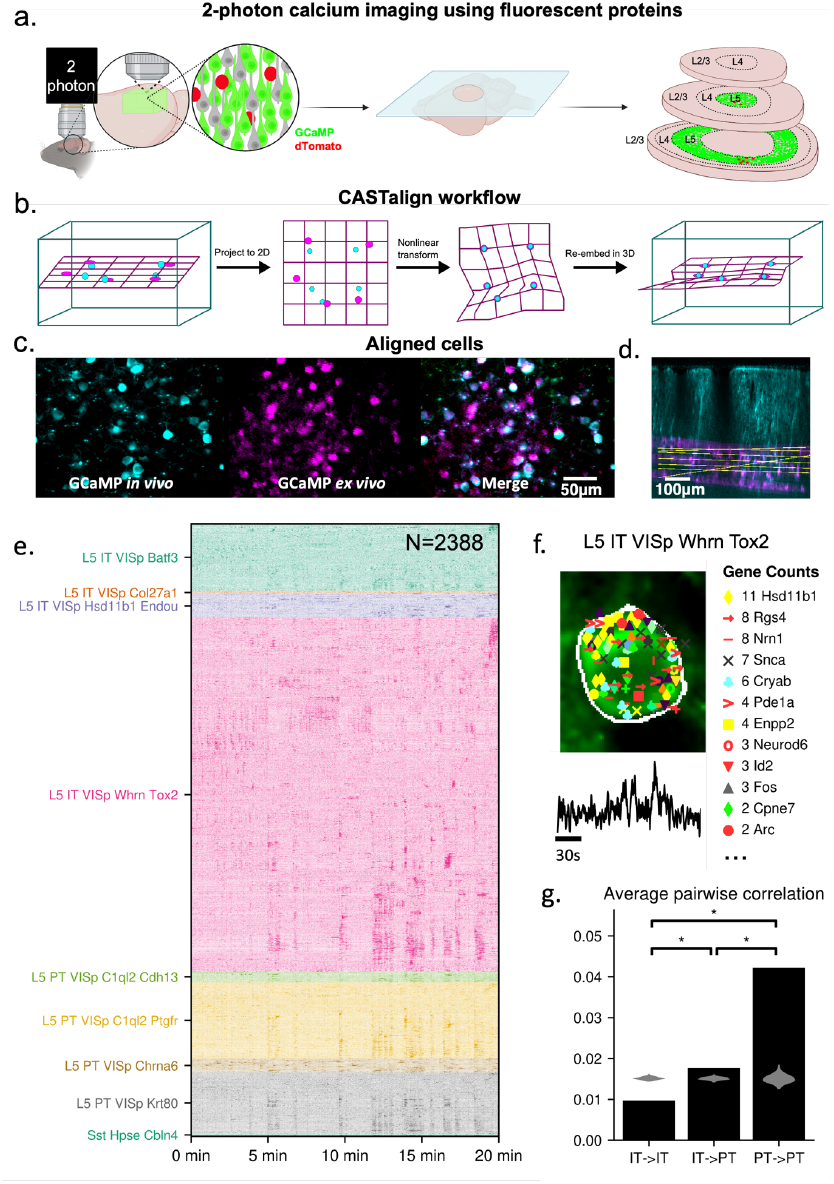
Registration of *in vivo* two-photon recordings to *ex vivo* coppaFISH 3D cell identities. (a) Experimental design for registration: two-photon calcium imaging is performed in tissue containing a sparse fiducial fluorescent marker, followed by sectioning and *ex vivo* processing. Figure created with BioRender.com. (b) CASTalign workflow for *in vivo* to *ex vivo* registration, using an initial projection to 2D, nonlinear transformation, and re-embedding in 3D. (c) Example of matched field showing GCaMP in an imaging plane *in vivo*, fluorescent signal *ex vivo*, and the merged alignment in x,y. Data from mouse F. (d) Matched sections across depth. *Ex vivo* GCaMP (magenta) is aligned to the high resolution *in vivo* GCaMP z stack (cyan). Yellow lines represent the *in vivo* imaging planes. (e) Activity heat map for 2,388 simultaneously recorded and registered neurons, ordered by transcriptomic identity, illustrating spontaneous activity across major layer 5 excitatory subclasses. (f) Example registered neuron showing *ex vivo* image data, assigned transcriptomic type, detected gene counts, and its *in vivo* calcium trace. (g) Mean pairwise correlations within intratelencephalic (IT) neurons, between IT and pyramidal tract (PT) neurons, and within PT neurons, showing stronger correlations among PT neurons than among IT neurons. * indicates <.01, two-tailed Monte Carlo permutation test. Grey indicates the distribution of correlations with shuffled identities.

CASTalign operates by building a graph connecting all images to be registered. Each node in the graph represents a volumetric image and its associated coordinate space, and each edge represents a coordinate transform mapping positions between the coordinate spaces of the two images. In our application of mapping *in vivo* 2-photon imaging to *ex vivo* transcriptomics, nodes correspond to *in vivo* imaging planes, *in vivo* z-stacks, *ex vivo* fluorescent protein images, and *ex vivo* gene positions (Supplementary Figure 6). Composing coordinate transforms along the edges of the graph allows any pair of images across any stages of our experiment to be aligned. Transform edges are learned either directly from data, or with varying levels of human input, proofreading and supervision, controlled from a graphical interface console. This provides a scalable way to register multiple images from large datasets.

The critical step of registration involves learning a map between the *in vivo* reference z-stack and the *ex vivo* fluorescent protein image (Figure 6b). This step is challenging because the required map is nonlinear, due to tissue warping that may occur during histological processing. To find the map, the user first manually matches a small number of fiducial cells between the two images. The program then uses these to fit a nonlinear mapping between the two stacks (see Methods). This mapping can then be refined in a graphical interface by adding additional matched cells if needed. In practice, this requires between 10-25 manually matched cells and takes about 15 minutes per tile, with approximately 3 tiles per section necessary to cover a 900 x 900 µm two-photon field of view. Figure 6c shows an example matched region. A full description of the semi-automated matching procedure can be found in the Methods, and is demonstrated in Supplementary Video 1.

We demonstrate this process in visual cortex. The mouse expressed GCaMP6s transgenically in layer 5 excitatory neurons using an Rbp4-Cre line, and GCaMP6s and dTomato were sparsely co-expressed virally with a pan-neuronal promoter to both provide fiducial landmarks and record neurons outside the Rbp4-Cre labelled population. We recorded spontaneous activity from 6735 neurons in primary visual cortex while the mouse viewed a grey screen (Figure 6d), then performed coppaFISH 3D and *in vivo* to *ex vivo* registration. CASTalign paired 3022 of the recorded neurons with *ex vivo* segmented neurons, of which 2388 passed our quality control criteria for unambiguous cell type assignment (see Methods). Figure 6e shows the GCaMP image, segmentation, anchor RCPs, gene reads, and activity trace for an example cell.

With these paired measurements, we compared activity in the two main transcriptomic classes of layer 5 excitatory neurons, pyramidal tract (PT) and intratelencephalic (IT) neurons. The raster plot suggested that IT neurons were more functionally diverse than PT neurons (Figure 6d). Pairwise correlations confirmed this: PT neurons showed higher within-class correlations than IT neurons, and IT neurons were more correlated with PT neurons than with other IT neurons (Figure 6f). The same analysis in a second mouse produced the same result (Supplementary Figure 5). These findings illustrate how large recordings with cell type identity can reveal structured activity within neural populations.

## Discussion

We have presented coppaFISH 3D and CASTalign, a platform for combining spatially resolved transcriptomics with *in vivo* two-photon imaging at scale. coppaFISH 3D detects mRNA with subcellular resolution in thick tissue sections that can be fixed by perfusion and cut with a vibratome, unlike most methods that require cryostat sections no thicker than a single cell. Gene expression can be linked to protein expression, allowing us to identify cell types that are vulnerable to pathology in diseases such as Alzheimer’s. The platform is fully open for use and extension by the community, powered by open chemistry, commodity hardware, and open source software for gene calling, cell calling, and nonlinear *in vivo* to *ex vivo* registration.

Several features make coppaFISH 3D especially well-suited for high throughput co-registration to neuronal recordings. First, it can be performed on commodity hardware at approximately £250 per section, compared to up to £5000 per slide for commercial platforms such as MERSCOPE and Xenium. Because coppaFISH 3D uses 50 µm vibratome sections, fewer sections are needed to cover the same tissue volume, further increasing this cost difference. Second, coppaFISH 3D quenches imaging rounds by stripping unneeded probes instead of cleaving fluorophores^46^, so failed experiments on precious tissue can be repeated if there are experimental errors, data corruption, or hardware failure. Finally, CASTalign exploits the 3D structure of thick sections for semi-automated registration. Thick sections are less prone to distortion than thinner frozen cryostat sections, substantially reducing tissue warping.

Although CASTalign was designed for coppaFISH 3D, it could also be used with other transcriptomics platforms. Other imaging-based 3D transcriptomics platforms, such as MERFISH^46^, STARmap^27^, HCR-FISH^47^ and others^23,48^, could likely use CASTalign with minimal modifications. We have also used CASTalign to register 2D transcriptomics to *in vivo* activity, demonstrating flexibility across different systems.

Compatibility with both *in vivo* recordings and IF staining makes coppaFISH 3D well-suited to studies of how disease processes affect different cell types. Although we do not record activity from disease models in the present study, antibody staining is compatible with our *in vivo* registration protocols, allowing disease-associated protein phenotypes to be linked to the activity of molecularly-defined cells. More broadly, coppaFISH 3D and CASTalign provide a practical framework for linking transcriptomic identity, protein expression, and *in vivo* activity at scale.

## Acknowledgements

We thank Steve Tovey and other members of Cairn Research Ltd. for aiding the development and maintenance of the imaging set up; Alexander Becalick for assistance with software; Anjali Singh for nomenclature; Takaomi Saido (RIKEN) for the use of the MAPT^S305N;Int10+3^ x APP^NL-F^ mice; Charu Reddy and Bex Terry for mouse colony maintenance.

This work was supported by the Medical Research Council (grant number MR/V003402/1), Wellcome Trust (223144/Z/21/Z, 108726/Z/15/Z), ERC (101097874) and Simons Foundation (NC-GB-IBL-00002672-1, SFI-AN-NC-IBL-00010540-05), to K.D.H. M.S was funded by the European Molecular Biology Organization LTF 712-2021. C.A.M, P.C, M.B and K.E.D are funded by the UK Dementia Research Institute through UK DRI Ltd, principally funded by the Medical Research Council. M.B is also supported by funding from the Cure Alzheimer’s Fund.C.A.M. was also in part funded by Eisai Co. Ltd. S.B. was funded by the European Union’s Marie Sk…odowska-Curie programme (835489). Y.I. was funded by Wellcome Trust and Gatsby Charitable Foundation.

## Author contributions

These authors contributed equally: Isabelle Prankerd and Maxwell Shinn. The gene panels were selected and designed by I.P., S.B and K.D.H. with input from Y.I. and M.S. RCP production was optimised for thick tissue sections by I.P. with the help of Y.I., K.D.H., D.O., A.R., S.B and Z.Z. Optimisation of the sequential imaging setup and protocol was done by I.P., Z.Z., C.A.M., M.B., Y.I., D.N., P.V.M.C., A.R., M.S. and K.D.H. The gene calling software coppafisher was developed by P.C.S., R.T., J.A.M.D. and K.D.H. with input from M.S., I.P., C.A.M., M.B., S.B., Z.Z., P.V.M.C. and D.N. MAPT^S305N;Int10+3^ x APP^NL-F^ mice, equipment and expertise were provided by K.E.D. The cell calling software pciSeq was developed and adapted by D.N. and K.D.H with input from M.S., C.A.M., I.P., M.B., P.V.M.C. IF alignment was developed by I.P., C.A.M., M.S., P.C.S., R.T. and M.B. The *in vivo* alignment software castalign was developed by M.S. All experiments and data analysis were carried out by M.S., Z.Z. and I.P. The figures were made by I.P. and M.S. and I.P., M.S., P.C.S. and K.D.H. wrote the manuscript with input from all authors. All work was supervised by K.D.H.

## Competing interest statement

The authors declare no competing interests.

## Materials and Methods

### Summary of coppaFISH 3D and CASTalign

Our platform enables high throughput identification of gene expression in neural recordings. It consists of coppaFISH 3D for transcript and cell type detection using spatially resolved transcriptomics, as well as CASTalign for co-registration of *ex vivo* transcriptomics, *ex vivo* antibody stainings, and *in vivo* neural recordings.

Broadly, coppaFISH 3D consists of gene panel selection, tissue preparation, RCP production, sequential imaging, and gene and cell identification steps. Gene panel selection only needs to be performed once at the beginning of a series of many experiments. Tissue preparation largely follows standard RNase-free histology practices. The RCP production is based on similar chemistry to *in situ* sequencing^29^ and to coppaFISH 3D’s predecessor, coppaFISH^18^, with the addition of procedures to allow reagents to permeate into 50µm thick tissue sections. Genes are identified from the sequential images, and cell types from the identified genes. Optionally, this can be followed by IF antibody staining.

After performing coppaFISH 3D, we use CASTalign to align the transcriptomics with IF and *in vivo* imaging. We perform functional and structural two-photon calcium imaging on the mouse prior to perfusion, and image the fluorescent proteins prior to sequential imaging. After performing coppaFISH 3D, we perform registration by learning a series of linear and non-linear transforms to register the *in vivo* imaging to the *ex vivo* imaged fluorescent proteins. Finally, we match the registered segmented *ex vivo* neurons with their corresponding *in vivo* ROIs.

All experiments were performed according to the UK Animals Scientific Procedures Act (1986).

### Gene panel selection and probe design

Before any experiments can be performed, a gene panel must be selected and DNA probes designed. coppaFISH 3D requires three types of oligonucleotide probe to be designed for each target gene. First, specific primers to enable the synthesis of complementary DNA (cDNA) through reverse transcription. Second, padlock probes to link cDNA sequences to gene-specific barcodes. Third, bridge probes to bind gene-specific barcodes and the specified dye probes. These three groups are pure DNA; in addition the method requires 7 dye probes (each being one fixed 20-nt sequence covalently linked to a fluorophore), and the anchor probe (a single fixed 20-nt sequence coupled to a fluorophore). These dye-coupled probes are much more expensive than pure DNA, but costs are manageable because only 8 are used.

Several aspects must be considered when designing a gene panel. First, genes may be of interest because they are used for cell typing, or because they are of some other biological interest. Among those used for cell typing, it is best to choose transcripts that are strongly expressed (as assessed in scRNA-seq data) but only in a subset of cells. Genes that are expressed in all cell types but in different frequencies often provide little information and can hinder the detection of more informative genes. This can pose problems not only for cell typing, but also for genes chosen for other biological interest.

We have found that a very simple approach to gene selection gives good results. This method is based on scRNA-seq data, and uses only scRNA-seq gene counts and not cell-type cluster assignments, which makes it invariant to cell type assignments. It is based on the principle that, because RNA levels of expressed genes are highly variable, there is much more information in the absolute presence or absence of a gene than its exact expression level. Thus, the best genes are those that are expressed strongly enough to be detected robustly in some cells, but completely absent in other cells. Specifically, for each gene we compute two numbers:

- ***X***_**99**_(***g***) is the 99th percentile of expression levels of gene ***g*** across the entire scRNA-seq database. This measures the expression level of gene ***g*** in cells of the classes that express it most, although it does not require explicit cell type assignments. The number 99 is selected to avoid the possibility of erroneously large counts in a small number of cells and can be increased to 99.9 or more if a large scRNA-seq database is used or if rare cell types are of interest.
- Second, ***Z***(***g***) is the fraction of cells whose expression of gene ***g*** is zero (for scRNA-seq data with UMI-based integer gene counts), or below some small threshold (for floating-point fPKM-based data; a suitable threshold can be found by examining histograms of expression levels and finding the edge of the peak at 0). A gene for which ***Z***(***g***) is 0.9 is a gene which is only expressed at all in 10% of cells; thus, expression of this gene signifies a low chance that the cell expressing it belongs to classes representing 90% of the total population.

Once both numbers have been computed for each gene, we select genes for which both ***X***_**99**_(***g***) and ***Z***(***g***) are large. Typically, we do this by making a scatter plot of the two variables with one point per gene, and by selecting all genes that a threshold for both numbers. The threshold is typically 0.3 for ***Z***(***g***) and for ***X***_**99**_(***g***) determined by ranking the genes passing the ***Z***(***g***) threshold and selecting a cutoff matching the total number of genes desired in the panel. Note that this procedure may undersample genes of low expression levels that nevertheless help identify some cell types; this can be corrected by manual verification that all types have sufficient genes and going further down the list if not.

The primer, padlock and bridge probe design process is described in the methods of Bugeon et al., 2022^18^. Padlocks are 80 nucleotides (nt), consisting of two 20nt cDNA binding arms, one 20nt barcode unique to the gene (but common to multiple probes against the same gene), and one 20nt anchor sequence (common to all genes). Padlocks are 5’ phosphorylated and ordered as 4nmol Ultramer plates (IDT). Primers are 15 nt (specific to where each padlock will bind) and 40 nt bridge probes (containing 20nt complementary to the gene barcode and 20nt complementary to the dye probe barcode) are ordered as 25nmol DNA plates (IDT).

### Tissue preparation

Mice are anaesthetised with isofluorane before being given a lethal intra-peritoneal injection of sodium pentobarbital (0.01 ml g−1). They are then transcardially perfused, first with RNase-free phosphate buffered saline (PBS, Gibco), then 4%(w/v) PFA (Electron Microscopy Sciences) in PBS until stiff. The brains are dissected from the skull, with great care to avoid damaging the tissue. RNase zap (Invitrogen) is used to clean the tools before and during extraction. The extracted brains are postfixed overnight in 4%(w/v) PFA in RNase free PBS at 4°C.

After post-fixation, brains are rinsed in RNase-free PBS and sectioned on a vibratome. Brains are glued to the stage and bathed in TE buffer pH8 (IDT). The brain is cut into 50µm sections (horizontally if aligning with in vivo data or any orientation for coppaFISH 3D alone) and collected in 24-well non-coated plates filled with freshly made sodium citrate buffer (1.5% W/V sodium citrate in distilled H2O and adjusted to pH 7, Sigma Aldrich). Once all tissue is collected, it is permeabilised by submerging the whole plate in a 70°C water bath for 30mins, which is sufficient for 50 µm tissue sections and only minimally bleaches native fluorescent proteins (which are required for registration to in vivo imaging). If no fluorescent proteins are present in the tissue, we instead set the water bath to 80°C which slightly increases the number of RCPs detected but bleaches fluorescent proteins. After water bath, excess buffer is removed and sections dried for 20mins before storage in a −80°C freezer. We have successfully performed coppaFISH 3D on sections that have been in the freezer for 3 years.

### RCP production

RCP production is the process by which mRNAs from our genes of interest are identified, bound by a probe with a unique gene barcode sequence and amplified. coppaFISH 3D uses free floating tissue sections (not coverslip mounted) to ensure reagent penetration in 50µm sections. The tissue is removed from −80°C and RNase-free PBS is added to the wells containing the sections of interest, allowing the sections to be easily picked up with a paintbrush. Sections are placed in a slide-mounted, un-coated 8-well plate (Ibidi) for RCP generation.

First, the sections are incubated in 200µl of reverse transcription mix at 42°C for 4 hours on a rocker. The reaction mix contains 0.5 mM dNTP mix (ThermoFisher), gene-specific primers (10nM each, IDT), 0.2µg.µl−1 Ul-traPure BSA (Invitrogen), 5mM DTT (Invitrogen), 1 U.µl−1 RNASE OUT RNase Inhibitor (Invitrogen) and 20U.µl−1 SSIV reverse transcriptase (Invitrogen) in 1x reverse transcription buffer (Invitrogen). After incubation the mix is fully removed and newly synthesised cDNAs are fixed in 4%(w/v) PFA in PBS for 30mins at room temperature (with no wash steps in between). The sections are then washed twice with PBS after fixation.

Next, padlock probe hybridisation and ligation are performed by adding 200µl of hybridisation mix. The hybridisation mix contains: 0.05M KCl (Sigma Aldrich), 20% ethylene carbonate (Sigma Aldrich), 1nM of each padlock probe (IDT), 0.2 µg.µl−1 UltraPure BSA, 0.3U.µl−1 Tth DNA Ligase (Qiagen) and 0.4U.µl−1 RNase H (NEB) in 1x Ampligase buffer (CamBio). Due to the different optimal temperature of the ligase and RNase enzymes, sections are incubated at 37°C for 1 hour and moved to 2-hour incubation at 45°C. After hybridisation, sections are washed twice in PBS.

Finally, rolling circle amplification (RCA) is performed by incubating in 200µl RCA mix overnight at 30°C. The RCA mix consists of 0.25nM dNTP mix, 0.2µg.µl−1 UltraPure BSA, 1mM DTT, 0.2U.µl−1 EquiPhi29 DNA Polymerase (ThermoFisher) and 1x EquiPhi29 buffer (ThermoFisher). Sections are washed twice with PBS and can be stored in PBS for approximately 2 weeks before sequential imaging. Longer storage results in the loss of signal when imaging. Samples with high autofluorescence may require the removal of background fluorescence to improve the signal to noise ratio during imaging. We found using a photobleaching lamp (Vizgen) for 11.5 hours at 4°C to be effective.

### Automated sequential 3D imaging

The automated fluidics system and the imaging procedure follows the protocol outlined in the methods of Bugeon et al., 2022^18^, while the imaging set up has been optimised for 50µm samples.

Images are acquired using an Eclipse Ti2E inverted microscope (Nikon) with a CFI Plan Apochromat Lambda 25x silicone immersion objective (NA 1.05, Nikon) which we chose as it combines a large field of view with high resolution, however other objectives will also be compatible with the system. An NIR-LDI 7 laser panel (89North) is used for illumination with the following wavelengths and nominal powers: 405nm (500mW), 445nm (1000mW), 470nm (1000mW), 520nm (500mW), 555nm (1000mW), 640nm (700mW), 730nm (900mW). An X-light V2 spinning disk unit (Crest Optics) with a 50-250 spiral disk is used to enable confocal imaging. This disk permits more light to reach the sample at the cost of some confocality, which allows us to use weaker dyes that would not be visible with a standard disk. Images can be captured by a single camera system with a filter wheel, but we obtained a 2-fold speedup in imaging using a quad cam setup (Supplementary Figure 2). The speedup is largely achieved by eliminating the time needed to move a mechanical filter wheel. Our setup consisted of 4 Orca-Fusion sCMOS cameras (Hamamatsu) connected via a series of dichroic beamsplitters, and fixed emission filters mounted in front of each camera at 400-480nm, 495-560nm, 565-647nm and a long pass filter of 655nm+. Diagrams of the imaging setup and 4-camera system can be found in Supplementary Figure 2. We use NIS elements (v.5.20.02, build 1453, Nikon) to select our region of interest (ROI) and acquire the images of multiple tiles, lasers and z planes using the ND sequence acquisition and multi-cam modules.

Tissue sections are mounted on coated coverslips, for which circular 40mm coverslips (Bioptechs) are dipped for 30 seconds in a solution of 4% w/v porcine gelatin (Sigma Aldrich) and 0.4% w/v chromium (III) potassium sulphate dodecahydrate (Sigma Aldrich) in distilled water. The coverslips are then dried in a 45°C incubator and stored in a dark box for up to 1 month. Sections are mounted using a paintbrush and dried for 2 hours at room temperature and fixed with 4% (w/v) PFA in PBS for 10 minutes. Samples for in vivo alignment are DAPI (ThermoFisher) stained during the fluorescent protein imaging round, while other samples should be manually stained with 10mM DAPI for 5 minutes.

Once the sections are mounted on the coverslip, we assembled the flow cell (FSC2, Bioptechs) and connected it to the automated fluidics system using two EFTE tubes (1/16-inch Red Delrin, IDEx Health and Science LLC). The fluidics system and automated chemistry protocol are described in Bugeon et al., 202218. The only difference is coppaFISH 3D requires a concentration of 20nM of each of the seven dyes: Atto425, AF488, DY520XL, AF532, AF594, AF647, AF750 and 500nM of DAPI in the dye mix.

First, we perform the fluorescent protein imaging round. This step can be skipped if not aligning to in vivo imaging. We image proteins in imaging buffer within the flow cell at 405nm, 470nm, and 555nm. Then, we disconnect the flow cell and remove the coverslip. We bleach using 5% SDS for 30 minutes followed by formamide for 30 minutes and then check the expression of the fluorescent proteins. If fluorescence remains, we continue bleaching with 30 minutes of 5% SDS and 1 hour of formamide until the section is fully bleached, checking after each round of chemical bleaching. Once fluorescent proteins are quenched, we reassemble the flow cell and continue with sequential imaging. We try to perform the fluorescent protein round and sequential imaging in fields of view that overlap as much as possible.

The imaging sequence acquires multiple tiles, z depths, and excitation wavelengths to cover the full ROI. Images are acquired in 16-bits in tiles of 2304 x 2304 pixels with 10% overlap. The full volume is captured in approximately 50-80 z-planes with a step of 0.9µm for each of the seven excitation wavelengths. The percentage power and exposure times used for each wavelength is as follows: 405nm (50%, 200ms), 445nm (80%, 300ms), 470nm (80%, 200ms), 520nm (80%, 300ms), 555nm (80%, 200ms), 640nm (80%, 200ms) and 730nm (80%, 200ms). The Nikon perfect focus system (PFS) is used to maintain focus over the imaging rounds and is set using the middle z-plane as a reference.

### Immunofluorescence staining

In samples where transcript detection is combined with IF, the immunostaining is performed after sequential imaging is complete. Mounted sections are removed from the flow cell, with the orientation marked on the coverslip to ensure it can be reassembled with a similar rotation. A hydrophobic barrier is drawn around the section using a pap pen (Vector Labs) and allowed to dry. Sections are washed twice on the coverslip with 2x SSC and can be stored in PBS for up to four weeks before staining.

The example section from a MAPTS305N;Int10+3 x APPNL-F mouse42 in Figure 5 underwent two rounds of immunostaining with 6E10 (biotin, mouse, 1:500, BioLegend), GFAP (chicken, 1:1000, Abcam), IBA1 (rabbit, 1:1000, Abcam) and P2RY12 (rabbit, 1:1000, Abcam). Each round consisted of blocking, primary incubation, secondary incubation and imaging. Between each round the section was stripped with a 10 minute incubation in the 4i elution buffer described in Hsu et al., 202443.

As 6E10 is a mouse antibody used in mouse tissue it requires more extensive blocking so the Mouse-on-Mouse immunodetection kit (M.O.M., Vectorlabs) and staining protocol was used. Blocking requires incubation for an hour at room temperature, after which the section is washed 3x in PBS before primary incubation for 48 hours at 4°C. Sections are then washed 3x for 5 mins, placed in a petri dish, covered with PBS and photo-bleached for 2-3 hours to remove any lingering fluorescence. Sections are then stained with secondary antibodies (ThermoFisher) at 1:400 concentration for 2 hours at room temperature in the dark. 10mM DAPI is added to the section in the last 15 minutes of the incubation. Sections are washed 3x for 5 mins, the coverslip is re-mounted, and the flow cell filled with imaging buffer. The section is manually coarsely aligned to the anchor image and imaged. The laser power and exposure time required is modified in an antibody specific manner to ensure a strong but not oversaturated signal.

### Gene identification

The process of identifying genes from the raw image data was performed by an open source Python package “coppafisher”, which is available at https://github.com/paulshuker/coppafisher.

The gene identification pipeline is divided into several independent stages. First, we Deblur microscope images, perform an Initial spot detection, then images are aligned through Registration. Then separate microscope tiles are Stitched together to create a single large image, and Initial spot calling takes a first estimate of gene identity and uses this to identify how the different fluorescent dyes bleed through into multiple channels. Finally, individual image pixels are optionally assigned to genes through Orthogonal Matching Pursuit (OMP) to disentangle overlapping genes, and a gene spot confidence score is computed.

#### Image Input

After sequential imaging, each set of images contains ***N***_***t***_ ≥ **1** tiles, each of which is a cuboid containing ***n***_***y***_ × ***n***_***x***_ × ***n***_***z***_ voxels, where ***n***_***y***_, ***n***_***x***_, ***n***_***z***_ are the number of pixels along the y, x, and z axes respectively (For the data shown here, the values were 2304 in the y and x dimension, and z ranges from 50 to 80). The voxel spacing of the ***x*** and ***y*** planes is equal, while the ***z*** spacing is typically larger. Each tile was imaged for ***N***_***r***_ = **7** combinatorial imaging rounds plus 1 anchor round. Adjacent tiles align along the ***x*** and ***y*** axes and have an approximate, known overlap percentage of 10%. On each round, every gene is labelled to one of ***N***_***d***_ = **7** dyes, according to a coding array of size ***N***_***g***_ × ***N***_***r***_ × ***N***_***d***_ which contains the Reed-Solomon codes assigned in the probe design phase.

Each sequence round uses ***N***_***c***_ = **9** colour channels (different excitation/illumination combinations from the 28 possible). The ***N***_***d***_ = **7** dyes each have a different spectral profile across these 9 colour channels; because the dyes are not perfectly spectrally separated and include long Stokes-shift dyes, the fluorescence profiles are not fully discrete, and are learned in a ***N***_***c***_ × ***N***_***d***_ “bleed matrix” as described later. Every sequence round also contains a channel with DAPI staining, which is spectrally distinct from all dyes (we found that blue dyes such as AF405 did not provide good depth imaging in confocal mode, so nothing is lost by allocating the short-wavelength channel to DAPI).

After all 7 combinatorial rounds, every imaging session also includes an anchor round, to which all rounds are later registered to. The anchor round uses the AF750 dye alone, to provide a high Signal-to-Noise Ratio (SNR) fluorescence at every RCP. Like all other rounds, the anchor round has a DAPI channel.

#### Devignetting and Debluring

The first stage of processing is to deconvolve all image stacks to compensate for the optics of the confocal microscope. To compensate for vignetting (the fading of intensities towards the edges of the images), we first divide each pixel’s value by an amount calculated as the median intensity at the same radius from the image centre (the values used can be found in supplementary material).

We then compensate for the microscope’s point-spread function using a Wiener filter. The filter uses an estimate of the microscope’s 3-dimensional PSF ***P***_***yxz***_ determined from symmetrised averages of high-quality spot fluorescence (the values used for ***P***_***yxz***_ can be found in supplementary data). The PSF is normalized

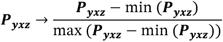

and then smoothly tapered by multiplying by the Hanning window49 of the same shape. Every image is then filtered using scikit-image’s wiener function50 with a regularisation parameter of **50**.

The result of the Wiener deconvolution is an image stack ***I***_***trcyxz***_ For every tile (***t***), round (***r***), and dye channel (***c***) and 3d image coordinate (yxz), stored as 16-bit floating point. Because of the deconvolution, the mean of this stack is close to 0 in each colour channel, nevertheless, its amplitude (standard deviation) can vary substantially between channels due to factors such as camera calibration.

#### Initial RCP production

The second step of the pipeline is to detect unambiguous gene detections: clear, bright spots in the image, which will be used to align images across rounds and channels later.

For every tile (***t***), round (***r***), and colour channel (***c***), a threshold is computed from the image stack 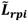 where ***z***_mid_ = floor(***n***_***z***_/**2**), |… | is the element-wise absolute value, and ***M***_***f***_ is a threshold multiplier set to **20**.

As each physical RCP will lead to several neighbouring voxels above threshold, we follow thresholding with a local maximum operation, keeping a voxel only if it is the brightest voxel in an ellipsoid with ***x***/***y*** semi-major axes of **5** voxels and a z semi-minor axis of **2** voxels.

#### Registration

The goal of registration is to register all images for one tile to a single coordinate system. The end product of registration is a look-up function which takes as input a coordinate (x,y,z), a tile number ***t***, a round ***r*** and colour channel ***c***, and returns the fluorescence of the appropriate image as registered to a common tile coordinate system. The tile coordinate system used is defined by the anchor image (the AF750 channel image on the anchor round, which labels every spot). In order to do this, the lookup function applies two coordinate transforms:

- A round-dependent nonlinear warping, to correct for tissue warping during processing and imaging, learned by optical flow registration of the anchor round DAPI image to each sequencing round’s DAPI image.
- An affine transformation from the AF750 image in each round, to every other colour channel in that round, learned via point cloud registration.

#### Optical Flow Registration

The aim of optical flow registration is to find a flow field ***L***_***rpi***_ containing the shift required for the ***i***th axis (i.e. one of the 3 dimensions) to match each voxel ***p*** = (***x, y, z***) on each sequencing round ***r***, to the corresponding voxel on the anchor round. This is done using the DAPI channel, which is present in all rounds and provides sufficient spatial resolution, following a global shift that is learned by phase correlation. The computation occurs in six stages, and is performed separately for each tile.

First, all images are downsampled by a factor of **4, 4, 1** along the ***y, x***, and ***z*** axes respectively, by taking the mean of non-overlapping blocks. This reducing the image sizes by a factor of16.

Second, the DAPI channel on every sequence round is rigidly registered with the DAPI image in the anchor round using an integer voxel shift, found via phase cross correlation.

Third, images are blurred with a Gaussian filter (standard deviation of **4, 4, 1** along the y, x, and z axes respectively.

Fourth, we find an initial raw flow field estimate 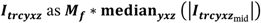 by the iterative Lucas–Kanade method (“optical flow”) method51. We use scikit-image’s function optical_flow_ilk with a window radius of **8** and prefilter set to true. To reduce compute time, the algorithm is run separately in a 5×5 grid of x-y chunks with 25% overlap, each spanning all z planes.

Fifth, we merge the flow fields found in each chunk into one global flow field for the entire tile. No interpolation is performed; where tile pixels lie on two overlapping chunks, the value at that pixel is taken from the chunk whose centre is nearest. Then, the combined flow field is upsampled to the initial voxel count by linear interpolation.

Sixth, we denoise the flow field based on accuracy. The Lucas-Kanade algorithm is unreliable in low-contrast regions. To overcome this problem, we identify regions where the flow field did, and did not, produce a good match of the round’s DAPI image and that of the anchor round, and interpolate flow fields from the former to the latter. In more detail, we expect Lucas-Kanade to work in regions where the original image has high but not low pixel intensity. We compute a score ***C***_***rp***_ = ***A***_***p***_***B***_***rp***_, for each voxel, ***p***, as the voxel-wise product between the anchor-round DAPI image ***A***_***p***_ and the flow-shifted DAPI image on sequencing round ***r B***_***rp***_. The product is then transformed 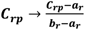, where ***a***_***r***_ and ***b***_***r***_ are the **25**th and **99**th percentiles of ***C***_***r***_ respectively, then clipped between **0** and **1**. Finally, the denoised flow field ***L*** is computed as a smoothed average of the raw flow field 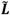, weighted by the product values:

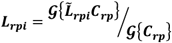

where 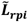 is the optical flow shift for the ***i***^th^ axis, and **𝒢**{…} is a three-dimensional Gaussian blur operator with standard deviation **10, 10, 5** along the ***y, x***, and ***z*** axes respectively. This operation therefore uses the values of 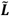 where ***C*** is large, which means the registration worked, to impute values of ***F*** where ***C*** is small and the registration did not work.

#### Iterative Closest Point

Once optical flow registration has produced a non-rigid alignment across rounds, this is further corrected by affine registration using Iterative Closest Point (ICP) registration52 based on the initial spot detection stage results. This compensates for factors such as chromatic aberration and other small shifts, scales, and rotations, that may differ between colour channels, rounds, and tiles. To reduce the number of parameters that must be estimated, for each tile we first learn a round-specific transformation then a channel-specific transformation. Thus, the total number of transformations learned is ***N***_***t***_(***N***_***r***_ + ***N***_***c***_) rather than ***N***_***t***_***N***_***r***_***N***_***c***_.

ICP finds an affine transformation between two point clouds, which we will call the source and target. The affine transformation is initialised as the identity. On each iteration, a spot from the source point cloud is matched to a spot from the target if the transformed source channel coordinate is within a radius of 5 voxels along x/y and 2 voxels along z of the target channel coordinate. Iterations proceed until the matched pairs do not change, up to a maximum of 50 iterations total. ICP is run independently for every tile.

To learn the round-specific transform, we first apply the optical flow corrections to the anchor image of the anchor round (the “anchor image”, which was made using the AF750 dye) on each tile, to transform it into a coordinate system for each round. We then apply ICP to find an affine transform using the nonlinear-warped anchor image point cloud as source, and each sequencing round’s image in AF750 channel as target. Note that the anchor image will have one spot for every RCP, but the AF750 channel in each sequencing round will have around ***N***_***d***_ times fewer; nevertheless, this does not present a problem since ICP only uses matched points to estimate the affine transform. This step thus learns ***N***_***t***_***N***_***r***_ affine transformations: one for each tile and round.

To learn the channel-specific transform. We run ICP again using as source the anchor image points transformed by both the nonlinear optical flow transformation and the round-specific transformation, and as target the detected spots in each colour channel separately. This step thus learns ***N***_***t***_***N***_***c***_ affine transformations: one for each tile and channel.

By composing the nonlinear flow from the anchor image to the AF750 channel in each sequencing round, then the affine transformations for each round and channel, we obtain a registration of all rounds and channels to a common tile coordinate system: given a coordinate in the anchor image, this provides a subpixel coordinate in each sequencing round and channel that matches it. Note that, to save memory, we do not explicitly construct transformed images, but instead rely on this transformation method to generate a subpixel-resolution coordinate in each tile, round, and channel for every pixel in the tile coordinate system, which is then used to retrieve fluorescence values by bilinear interpolation.

#### Stitch

Next, the tile coordinate systems are all aligned to a single global coordinate system for the entire tissue section. Again, we do not explicitly stitch images together to save memory, the stitching takes place at the level of coordinate systems.

The stitching transformations are learned using the anchor round DAPI images, making use of the 10% overlap between tiles. First, the anchor round DAPI images for each tile are multiplied by a Hanning window of the same shape. Then, for tile t and its i^th^ adjacent tile, a three-dimensional shift vti, is calculated by sub-voxel phase cross correlation^53^ on the tile-overlap region. Every shift has a corresponding reliability score, λti, that is the square of the Pearson correlation coefficient between images after the shift vti is applied.

A tile has up to four shifts, computed relative to each of its neighbours. Our aim is to produce a global coordinate system, which means a single shift ***S***_***t***_ per tile; this problem is overdetermined. To gather one shift per tile, we define a loss function that weights the similar every shift weighted by its score λti such that the loss function

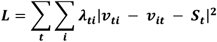

is minimised to calculate the final tile shifts, ***S***_***t***_. Because this is quadratic in ***S***_***t***_, it can be solved by standard least-squares fitting.

#### Initial Spot Calling

Spot-calling (i.e. identifying the genes underlying each RCP) is based on a model for how the ***N***_***r***_***N***_***c***_-dimensional sequencing images depend on the underlying genes. In principle, we can predict this by summing fluorescence vectors for every gene using the designed codes. In practice however, the actual images often deviate from the theoretical predictions. For example, inconsistencies in DNA probe manufacture could result in bridge probes for some genes in some rounds occurring at lower concentrations than expected, leading to a deviation of the actual fluorescence vectors from the theoretical prediction. Alternatively, fluctuations in camera gain or shutter dynamics could lead to variations in the global image brightness that are specific to certain tiles, rounds, and channels.

To compensate for these deviations, we perform spot calling in two phases. The first phase (“initial spot calling”) aims to estimate parameters that quantify these deviations. This phase does not require resolution of overlapping spots, so focuses only on the well-isolated spots detected in initial spot detection. The second phase (“orthogonal matching pursuit”) uses these parameters to produce a dense gene detection, resolving overlapping spots assuming linear addition of brightnesses from each spot.

#### Initial Normalisation

Let ***F***_***src***_ denote the intensity in colour channel ***c*** and round ***r*** of spot ***s*** detected by initial spot detection; to create the array ***F***_***src***_ we use the spot locations detected on the anchor image by initial spot detection as input to the fluorescence lookup function, that applies the learned registrations and retrieves a ***N***_***r***_***N***_***c***_-dimensional fluorescence vector for each spot. ***F***_***src***_ is divisively normalised separately for each tile, round, and channel, assigning a value of 1 to the 95^th^ percentile of fluorescence values in that tile round and channel.

#### Initial Gene Assignment

Next, we find an approximate gene assignment for a subset of high-quality spots, in a manner that does not require accurate values for the parameters. To do so, we assume that the L2-normalised intensity vector of spot ***s***, de-fined as 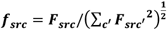, comes from a Fisher-von Mises distribution whose mean ***b***_***grc***_ depends on the actual gene ***g***. The Fisher-von Mises distribution is a distribution on the sphere, whose probability density is:

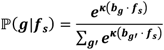

The preliminary vectors ***b***_***g***_ are ***N***_***r***_***N***_***c***_-dimensional preliminary code vectors for each gene ***g***, given by 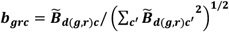. Here, ***k*** = **3** denotes a concentration parameter, and ***d***(***g, r***) denotes the dye assigned to gene ***g*** in round ***r*** during probe design. 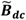 denotes a preliminary bleed matrix (for values see supplementary data) giving the spectral intensity of dye ***d*** in colour channel ***c***, obtained by imaging free-floating drops of dye, but found not to be perfectly accurate, requiring later correction by the algorithm.

The assignments of spots to genes made by Fisher-von Mises assumption will miss many genuinely detectable spots. However, because these spots are only used to estimate parameters such as scaling, this does not present a problem. Only well-classified spots with probability ℙ(***g***|***f***_***s***_) ≥ **0. 9** for the best ***g*** are retained for further analysis.

#### Bleed Matrix Calculation

The bleed matrix ***B*** estimates the amount of fluorescence expected in channel ***c*** from dye ***d*** (it has this name due to “spectral bleeding” of dyes appearing in multiple colour channels). To estimate it, ***N***_***c***_-dimensional colour vectors representing dye ***d*** are gathered from all spots and rounds which are expected to use dye ***d*** in that round. We then construct a ***N***_***c***_-dimensional spectral vector for dye ***d*** as the largest magnitude right singular unit vector of this matrix. We stack those vectors together for each dye to form the ***N***_***d***_ × ***N***_***c***_ size bleed matrix ***B***.

#### Bled Code Calculation

Our next task is to compensate for systematic deviations in the actual fluorescence intensity profile measured for each gene from that expected from the original probe design. If there were an equal concentration of each bridge probe for each gene, then we would expect the fluorescence of a spot of gene ***g*** in round ***r*** channel ***c*** to be ***B***_***d***(***g***,***r***),***c***_, purely determined by ***B***_***dc***_ of the dye ***d***(***g, r***) assigned to this gene in this round. We term the matrix ***B***_***d***(***g***,***r***),***c***_ the “raw bled code” as it takes into consideration the spectral bleed matrix ***B***.

However, in practice, the concentrations of manufactured bridge probes varies between genes and rounds, the fluorescence of some genes appears dim or bright in some rounds, in ways that are reliable across spots and tiles, but unpredictable *a priori*.

To compensate for this variation, we estimate an array ***E***_***grc***_ which contains the expected intensity of gene ***g*** in round ***r*** channel ***c***, which we term the “free bled codes” as they can deviate from the original bled codes. ***E***_***grc***_ is estimated by Bayesian inference, with a structured prior that easily allows scaling of the magnitude of the ***N***_***c***_-dimensional fluorescence vector of a gene in a round (which frequently occurs due to manufacturing inconsistencies), but strongly penalises deviations from the direction of this vector (which would require off-target binding and is observed rarely). The resulting formula is:

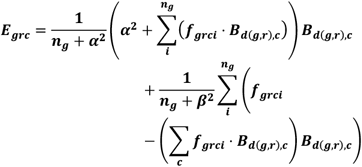

Here, ***n***_***g***_ is the number of spots assigned to gene ***g*** with probability ≥. **9**; ***f***_***grci***_ is the observed fluorescence in round ***r***, channel ***c*** of the ***i***^th^ spot assigned to gene ***g***; ***B***_***dc***_ is the bleed matrix calculated above; and ***α***^**2**^ = **10, *β***^**2**^ = **50** are parameters representing the prior precisions for variations along and orthogonal to the bled code vectors. These parameter values mean that approximately 10 spots of one gene must be observed before the overall scaling changes, but 50 spots must be observed before the relative brightness of different channels within a round is altered.

Round and Channel Normalisation. To compensate for experiment-to-experiment variability in factors such as imaging gains and dye probe concentrations, we learn a matrix ***V***_***rc***_ that scales the free bled codes ***E***_***grc***_ in a gene-independent manner. ***V***_***rc***_ is found by minimising the loss function

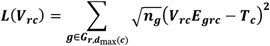

where ***d***_max_(***c***) is the dye that appears brightest in channel ***c*** (***d***_***max***_ = [**0, 1, 1, 3, 2, 4, 5, 5, 6**]). For each of the channels, ***T***_***c***_ is a target value that we intend to scale the mean value of spots to in that channel. We do not use the same value of ***T***_***c***_ for all ***c***, because we have more colour channels than dyes, which helps with dye detection, but means we downweight multiple channels that respond to a single dye. The specific values used for ***T***_***c***_ were [**1, 0. 8, 0. 2, 0. 9, 0. 6, 0. 8, 0. 3, 0. 7, 1**]. The factor 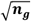 is used to avoid the estimate of this matrix being overly dominated by very strongly expressed genes.

This scaling factor allows us to construct the final gene bled codes ***K***_***grc***_, as a L2-normalised version of the product ***E***_***grc***_***V***_***rc***_:

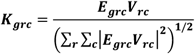

In addition to the entries of ***K***_***grc***_ for the genes in the gene panel, we add ***N***_***c***_ “background genes” whose fluorescence vectors are constant across rounds, and equal 1 for colour channel ***c*** in all rounds, for ***c*** = **1** … ***N***_***c***_. These background genes allow the algorithm to automatically detect background fluorescence from the native tissue, such as lipofuscin, and prevent it leading to erroneous gene detections.

#### Tile Normalisation

A second form of correction factor is learned to compensate for fluctuations in laser power or shutter opening, which can lead the intensity of different colour channels to vary between rounds, channels, and tiles in a manner that is consistent across all spots in a tile but unpredictable *a priori*. We will estimate these fluctuations by a scale factor ***Q***_***trc***_ measuring the deviations from nominal power for every image exposure, i.e. every tile ***t***, round ***r***, and channel ***c***, but independent of gene ***g***.

To do so, we first compute for each tile ***t*** an array ***D***_***tgrc***_, the same way as the array ***E***_***grc***_ defined above, but using only the set of spots assigned to tile ***t***. The scale factor ***Q***_***trc***_ is found for every tile ***t***, round ***r***, channel ***c*** by minimising the loss function

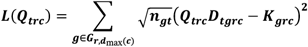

where ***n***_***gt***_ is the number of gene 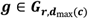 spots in tile ***t***. This scale factor therefore estimates the optimal tile scaling required to allow a single bled code matrix K (whose deviations from nominal depend on chemical factors that will be equal across tiles), to work on all tiles. The factor of 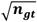 was chosen to prevent genes with many detections from dominating the estimate.

Finally, the full colour normalisation factor is constructed as ***A***_***trc***_ = ***Q***_***trc***_/***Y***_***trc***_, where ***Y***_***trc***_ is the 95^th^ percentile of the spot fluorescence values of round r channel c in tile t. Thus, multiplying the fluorescence values by ***A***_***trc***_ will scale all spots to the same intensity scale as the gene bled codes ***K***_***grc***_, thus allowing orthogonal matching pursuit to be used uniformly on all tiles.

#### Orthogonal Matching Pursuit

Once the scale factors have been learned from the initial spot calling, we rerun gene detection using a more sophisticated method, Orthogonal Matching Pursuit (OMP), which can resolve overlapping spots. OMP produces the final gene reads.

OMP works on the full voxel arrays, not just the locations of detected spots, and thus detects many more reads than initial spot calling. It uses the sequencing rounds and channels but not the anchor round and DAPI channels. The voxel values for all rounds and channels are registered using the transformations found in the Registration stage via bilinear interpolation. Then they are multiplied by the full colour normalisation factors, ***A***_***trc***_, calculated in Tile Normalisation stage for each tile/round/channel pair to get a normalised colour vector ***S***_***prc***_ for the ***p***^th^ voxel in round ***r***, channel ***c***. The aim of OMP is to find a sparse set of weights ***w***_***pg***_ summarising the amount of fluorescence detected in voxel ***p*** from gene ***g***, so

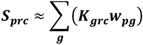

This is achieved by iterating two steps for each voxel ***p*** until all voxels reach a stopping criterion, that differs between voxels. Each iteration adds another gene to voxel ***p***, and terminates the loop for that voxel if no suitable genes are found. Thus, on each iteration ***i***, the algorithm is only applied to a non-increasing “active set” of voxels, starting with every voxel.

#### Step 1: Next gene assignment

If a voxel is active on the ***i***^th^ iteration then ***i*** − **1** genes have already assigned been it. Each assigned gene ***g*** will in the previous iteration ***i*** − **1** have been assigned a weight ***w***_***pg***;(***i*.1**)_. To see if another gene can be usefully fitted, we compute the latest residual voxel colour, ***R***_***prci***_ as

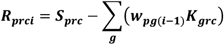

where ***K***_***grc***_ is the gene ***g***’s bled code in round ***r***, channel ***c***, normalised across rounds and channels so that ‖***K***‖_***g***.._ = **1** for all ***g***, where ‖***X***‖_…_ represents the L2 norm of ***X*** over all indices replaced by a dot (.). Note that ***R***_***prci***_ = ***S***_***prc***_ on the first iteration (***i*** = **1**).

The algorithm used is a modification of the standard orthogonal matching pursuit algorithm, adding a variance factor that accounts for the variability of fluorescence compared to gene bled codes. The reason for this modification is we have found that fluorescence vectors can differ substantially from one spot to the next, even of the same gene. Failing to account for this variability led to detection of spurious genes that attempted to compensate for deviations from the actual fluorescence vector of a gene ***g*** from the predicted vector ***K***_***g***_.

For every candidate gene 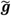 to add on iteration ***i***, a score is computed as

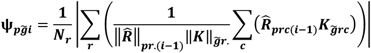

where

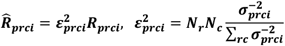

and

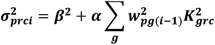

The factor 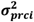 represents the amount of variance in the fluorescence expected in round r channel c voxel p, given the assignments made on iterations 1… i-1. The score 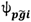 is proportional to the log-likelihood-ratio in predicting the observed fluorescence vector, that would be obtained by adding gene 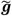 to voxel ***p*** on round ***i***, assuming a Gaussian distribution for fluorescence. The parameters took the values ***α*** = **120**, and ***β*** = **1**. Since ***α*** ≫ ***β***, the variance term **ε**_***prci***_ down-weights ***r*** / ***c*** pairs that are already accounted for by previous gene assignments.

After the scores are computed, gene 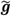 is assigned to voxel ***p*** in round ***i*** if all the following conditions are met:

- 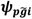 is the largest gene score of all genes.
- 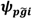 > **0. 5**.
- 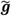is not already assigned.
- 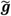 is not a background gene.
- The residual colour’s intensity min_***r***_ 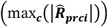 > minimum_intensity_***t***_.
- ***i*** ≤ **5**.

The intensity threshold minimum_intensity_***t***_is defined as minimum_intensity_***t***_= **4 *median***_***p***∈***middle z plane***_ (min_***r***_ (max_***c***_(|***S***_***prc***_|))) If the conditions are not met for any gene, then the voxel is no longer iterated on. In other words, iterations stop once a low-quality match is found, a match to background fluorescence occurs, the residual fluorescence is too dim, or 5 gene assignments have already been made.

#### Step 2: Gene weights

Once genes have been assigned, we update each the gene weights ***w***_***pgi***_ for all voxels with new genes by minimising the loss function

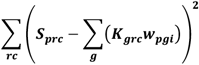

#### Spot Detection and Scoring

The result of OMP is a sparse assignment of up to 5 genes to each voxel, with weights accompanying each assignment. Detections of a single gene will tend to be spatially clustered around genuine gene locations, due to the physical size of the RCP and point spread function; occasional false assignments can be made but are less likely to be spatially clustered.

To turn these assignments into a discrete set of high-confidence gene reads, we compute scores quantifying the quality and confidence of each detected gene. First, every assigned gene is assigned a voxel score, ***c***_***pgi***_representing the decrease in log likelihood that would occur if that gene were excluded from the list of genes assigned to that voxel:

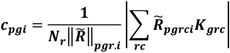

where

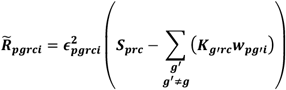

and

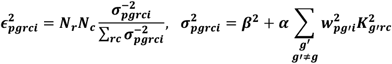

***c***_***pgi***_ is made negative if ***w***_***pgi***_ < **0**. For an unassigned gene 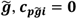.

#### Gene Spot Confidence Scores

To generate confidence scores for each gene detection, a spatial running mean of the 3D voxel scores array is computed by convolving with a 5×5×3 smoothing kernel (for values see supplementary data), normalizing by the result of convolving a matrix of all 1s with the same kernel, and padding with zeros to preserve the image size. Because a single RCP will result in multiple high-scoring neighbouring voxels, we finally apply the thresholding and local-maximum-detection algorithm described in the Initial spot detection section to the convolved gene scores, with a score threshold of **0. 1**, x/y radius of **3**, and z radius of **2** to find the final locations and spot confidence scores of all detected genes.

### Cell type identification

After genes have been identified, we classify each cell’s transcriptomic identity. We first identify all of the cells by performing segmentation on the 3D confocal images and then use a Bayesian classification algorithm to assign gene reads to segmented cells, using the assigned genes to determine the cell’s identity using an existing single cell RNA-seq taxonomy. We base our method on the Bayesian classification algorithm pciSeq^38^, with two main modifications. First, we modified the segmentation algorithm to improve 3D segmentation, and second, we modified the Bayesian algorithm to use 3D spatial information for gene assignment.

#### Segmentation

The data used for 3d segmentation depend on whether the transcriptomics is registered to in vivo imaging. If so, we segment the GCaMP image obtained ex vivo during the fluorescent protein imaging round, which provides a good outline of all imaged cells. This method identifies all neurons which underwent in vivo imaging, but since not all cells express GCaMP, this method only identifies a subset of all cells. Otherwise, we segment the DAPI images from the anchor round. Segmentation was performed using custom-trained Cellpose^37^ models, operating in 3D mode.

We found that Cellpose’s 3D mode frequently gave segmentation errors characterised by oversplitting and checkerboard-like cleavage planes. This is because Cellpose’s 3D mode assumes that all dimensions have equal imaging quality. However, the quality of our confocal images was higher in the x-y dimensions than in the z dimensions due to the use of non-square voxels and a non-spherical point-spread function.

To address this issue, we modified the Cellpose 3D algorithm to place more emphasis on segmentations computed from x-y slices. In the standard implementation, Cellpose processes each plane orientation (x-y, x-z, and y-z) independently: for every slice it estimates a 2D flow (the gradient pointing toward the cell centre) and a pixel-wise cell probability, producing three volumetric stacks of gradients and three stacks of probabilities. It then constructs the final 3D flow field by averaging the x, y, and z components from all three gradient stacks with equal weight and similarly forms the 3D cellprobability volume by averaging the three probability stacks. In our customised version, the x and y flow components and the voxel-wise cell probabilities are taken exclusively from the x-y estimates, where image quality is highest. To compute the z component, we use the average of the z components from the x-z and y-z flow estimates. As a result, high-quality x-y information drives nearly all aspects of the segmentation, while lower-quality orthogonal views contribute only to the z direction. This change substantially improved segmentation quality and yields 3D cell outlines for every cell in the volumetric confocal image. Our modifications are available at https://github.com/mwshinn/cellpose3dplus.

#### Cell calling

We used a probabilistic algorithm in conjunction with an existing scRNA-seq taxonomy to identify each neuron’s transcriptomic type. This algorithm is a modification of an existing probabilistic model (Qian et al., 2019^38^), which uses variational Bayesian approximation to assign each gene a probability of belonging to each of its neighbouring cells, as well as the probability of each cell belonging to each potential cell type. We modified this existing algorithm to operate on three-dimensional data. Transcript locations and segmented cell boundaries are represented in 3D and the spatial relationships between transcripts and candidate cells are computed in three dimensions rather than in 2D sections. The probabilistic framework remains unchanged but all spatial calculations and neighbourhood assignments use the full 3D geometry of the tissue volume. When analysing primary visual cortex only, we used the RNA sequencing data and taxonomy from Tasic et al. (2018)^40^, and when analysing the entire brain, we used Yao et al. (2023)^39^. Our cell assignment data can be viewed using our web-based viewer (https://acycliq.github.io/coppafish3D_hippocampus/. Our cell calling algorithm is available as the pciSeq Python package (https://github.com/acy-cliq/pciseq_3d).

### Two-photon calcium imaging

We perform in vivo two-photon calcium imaging on awake, behaving mice using mouse models and imaging protocols optimised for registration to ex vivo tissue sections. We use mice that express both a fluorescent calcium indicator as well as a sparse fiducial fluorescent protein marker to assist with registration. Mice were allowed to run on a treadmill during imaging and were surrounded by three screens to display visual stimuli. We performed imaging across at least two sessions: one session to image high-quality reference z-stack volumes for registration, and subsequent sessions with quick daily z-stacks paired with multi-plane recordings, all of which were registered to the reference z-stacks so that all imaging used a common coordinate system.

Registration of in vivo two-photon calcium recordings to ex vivo transcriptomics makes use of both a calcium indicator as well as a sparsely expressed fiducial marker to facilitate registration. The data analysed in this paper, was imaged in the primary visual cortex of a mouse that expressed GCaMP6s transgenically in excitatory neurons in layer 5. The mouse also had sparse stereotactic injections of a virus coexpressing GCaMP6s and dTomato, which both facilitated registration, and allowed GCaMP imaging from virally-infected inhibitory neurons. Two transgenic Ai94(tetO-GCaMP6s)-Camk2a-tTa-RBP4-Cre mice underwent stereotactic surgery to implant a circular 3mm cranial window over VISp at coordinates ML=-2.0mm, AP=-2.7mm. A head plate was implanted surrounding the cranial window to enable head-fixation, sealed with a glass cranial window, using cyanoacrylate adhesive (Vetbond, 3M) and dental cement. During this procedure, the mouse was stereotactically injected with 138nL of AAV-hSyn1-GCaMP6s-P2A-nls-dTomato (Catalogue #51084-AAV1, Addgene) at a depth of 300µm using a bevelled micropipette with a Nanoject II injector attached to a stereotaxic micromanipulator. After recovery, the mouse was habituated for handling and head-fixation for three days before carrying out recordings. In vivo recordings were preformed 25 days after the virus injection using a commercial two-photon microscope with a resonant-galvo scanhead (B-scope, ThorLabs) controlled by ScanImage.

During the first imaging session, we recorded two reference z-stacks which stretched from above the brain surface to the maximum visible depth, one in each the green and red channels. In order to sample across a large range of timepoints for neurons at all depths and capture as much fluorescent activity as possible, we imaged the z-stacks with a high-speed piezo objective scanner (“fast-z”) in three or four 300µm chunks each, which we subsequently stitched together. One z-stack was recorded with a 920nm and the other at 1060nm, both with 100 planes at 3µm spacing using both green (λ=525/50) and red (λ=607/70) emission filters in each image. Laser power and imaging time was adjusted for each of the chunks at the time of imaging to enhance signal brightness and reduce noise, with imaging time ranging from 5-20 minutes per chunk (90-360 repeats per voxel), and longer durations used for the depths that are targeted for recording. For the portions of the 920nm z-stack that overlapped with the imaged area, stack chunks were obtained while the mouse viewed a visually salient stimulus such as zebra noise (Skriabine and Shinn et al., 2026^54^) to increase the probability of detecting sparsely-firing neurons.

During subsequent imaging sessions, we recorded a quick daily z-stack at the beginning, which we would subsequently register these to the reference z-stack. This strategy provides the flexibility of aligning multiple imaging sessions from the same mouse to the same common reference coordinate system. These daily z-stacks were nearly identical to the reference z-stacks, except to minimise imaging time, we only collected a z-stack with λ=920 excitation wavelength, and each chunk was imaged for only 1-2 minutes (18-36 repeats per voxel). We then recorded calcium activity with 7 planes at 30um steps while the mouse looked at three grey screens surrounding it in a virtual reality setup, with a frame rate of 4.28 hz.

Two-photon imaging recordings of neuronal activity were processed using Suite2p. The result was deconvolved and reconvolved with a Gaussian filter with standard deviation 0.3s and interpolated to 10 hz, taking into account the differences in timing for neurons in different imaging planes.

Z-stacks were constructed from imaged chunks through stitching. First, the temporal average of the z-stack chunk acquisitions was computed to obtain a volume for each chunk. We did not adjust for movement, as the low signal-to-noise regimes with few neurons in the deepest parts of layer 5 produced unreliable motion-corrected registrations. Stitching was performed in the z dimension only by finding the z offset that maximised the correlation of the overlapping region of adjacent z-stack chunks. All results were checked manually, and when converted to microns, we confirmed that the computationally-determined offsets were identical to the offsets measured during imaging.

### CASTalign registration framework

We developed the CASTalign (CoppaFISH, Antibody Staining, and Two-photon alignment) framework to register large, heterogeneous 3D imaging datasets across modalities and experimental stages.

CASTalign is a general framework, designed for co-registration of hundreds of 3D images to a single coordinate system. It does this by building a graph in which each node represents a volumetric image, and each edge is a linear or nonlinear transformation aligning the two nodes connected by it. Once sufficient edges have been added to make the graph connected, i.e. there is a (potentially multistep) path between each pair of points, it is possible to align any image to any other image using chains of transformations.

In our application, the images correspond to the *in vivo* imaging planes, *in vivo* daily and reference z-stacks, *ex vivo* confocal imaging of GCaMP and dTomato proteins, and *ex vivo* transcriptomic gene detection (Supplementary Figure 6). CASTalign is also used to manually proofread the automatically-generated transforms joining microscope image tiles, and correct where necessary.

CASTalign consists of three main components:

#### Transforms

Given a pair of volumetric images, each representing a different coordinate space, a transform converts coordinates from one image’s coordinate space to the other’s. The transforms can be affine (for example to correct for camera rotation or translation) or nonlinear (for example to correct for tissue warping). Because transforms are performed analytically, and all transforms we used are invertible, they can be composed into long bidirectional sequential chains and performed losslessly in a single interpolation step, eliminating the need for downsampling and resampling.

#### Transform graph

When aligning many different images to each other in the same dataset, it can quickly become unwieldly to organise all of these transforms and their relationship to each other. In CASTalign, individual images are represented as nodes on a transform graph, and chains of transforms connect the nodes. This means that any points or images in one node’s coordinate system within the graph can be transformed to any other by chaining together the transforms of the edges in their shortest path. This is performed in one resampling step, avoiding the cumulative image degradation associated with repeated sequential transformation

#### Graphical interface

CASTalign has a graphical user interface (GUI) which can be used to create linear or nonlinear transforms, or to inspect automatically generated transforms and adjust if necessary. The GUI overlays the target image on top of the transformed source image by displaying them in different colours. CASTalign allows two different methods for specifying transforms in the GUI.

In “parametric mode”, the GUI allows the user to specify parametric transforms (which must be linear) by dragging the source image on-screen, and specifying additional parameters such as rotations, dilations, and shears using sliders and check boxes.

In “point pair mode”, the GUI allows the user to specific either linear or nonlinear transforms by clicking on pairs of matching points from the source and target images. The transform is updated in real-time as new point pairs are added. This shares similarities to BigWarp^55^, but while BigWarp is primarily focused on pairwise alignment of images, CASTalign is intended for general multi-stage, multi-image registration spanning many related datasets, including a mix of automated and manual registration.

For many transform steps, a first estimate of the transform is computed automatically, for example by phase correlation or point cloud registration. In all cases, these are proofread by a human operator in the GUI and corrected if necessary.

These features benefit both IF alignment as well as *in vivo* to *ex vivo* registration, where accurate correspondence needs to be maintained across large fields of view, multiple modalities, and several intermediate coordinate systems. CASTalign is available through the Python package castalign (https://github.com/mwshinn/castalign).

#### Types of transforms

Parametric transforms. All transforms specified by the parametric GUI mode must be affine: linear transformations plus shifts specified by a matrix ***A*** and shift ***b***, sending a vector ***x*** ↦ ***Ax*** + ***b***. Affine transforms include translations, dilations, rotations, and shears. To specify an affine transform in parametric mode, the user can drag the source image along the screen with the mouse, or enter rotation angles, dilations, shears, or matrix elements through sliders.

Point-based transforms. Transforms specified by the point-matching interface can be affine or non-linear. Once the user has clicked on a set of point pairs, the GUI automatically fits a transform from one of a list of transform families; which family to use for any edge is chosen by the user.

- **Translation:** A translation is fit by aligning the centroid of the source to the target image.
- **Rigid:** A rotation and translation are fit by aligning the centroids and finding the optimal rotation using orthogonal Procrustes analysis.
- **Affine:** Fit by linear regression.
- **Laminar affine transformation:** In tissue sections, the z dimension is much thinner than x and y, making rotations and shears in z difficult to estimate. Fortunately, these are uncommon: coverslip-mounted sections usually warp in x and y, but only expand or contract in z. To register these images, we therefore fit a constrained affine transform allowing rotation and shear within the x-y plane, but only scaling along the z direction (i.e. a 2,1 block-diagonal matrix). To find this transformation, CASTalign first rotates the demeaned points in the source and target images so that the direction of minimum variance is in the z direction (this rotation is found by singular value decomposition). It then fits two separate affine transformations: a 2D affine transformation in the x and y dimensions (ignoring the z dimension), and a 1D transformation in the z dimension (ignoring x and y), and finally generates the full transform by composing the source rotation, block affine transform, and inverse target rotation. This is given by **A** = **RM*R***^−**1**^ where R is the rotation matrix that puts the dimension of lowest variance in the z dimension, and M takes the 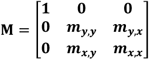.
- **Laminar triangulation:** When sample warping has occurred in coverslip-mounted slides, the laminar affine transformation is not sufficient. In such cases, the user can specify to instead use laminar triangulation. This transform consists of nonlinear warping in the x-y plane but only translation in the z dimension. To find it, CASTalign again rotates both point clouds such that the direction of minimum variance is in the z direction. It then defines a 2D Delaunay triangulation using the x and y coordinates of the rotated source point cloud, ignoring the z coordinate. For each triangle, it finds a constrained affine transform using a similar principle as the laminar affine transform. Specifically, each triangle within the triangulation is fit to the transformation matrix **A** = **RM*R***^−**1**^, where R is the rotation matrix that puts the dimension of lowest variance in the z dimension, and M takes the form 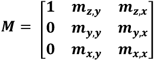. Finally it generates the full transform by composing the source rotation, triangle-wise affine transformation, and inverse target rotation. Points whose x-y coordinate is beyond the convex hull of the keypoints are transformed according to the triangle whose perimeter is closest.

#### Aligning IF with gene detection

To register immunofluorescence to coppaFISH 3D, we use the DAPI channel of both images. First, the raw IF tile images are stitched together. In overlapping areas, values are estimated via linear interpolation. We then register the stitched DAPI channel of the IF image to the stitched DAPI channel from the anchor round of the sequential imaging using a rigid registration. To fit this using CASTalign, an operator selects about four DAPI nuclei that can be identified in both the IF image and the sequential imaging (about 5 minutes per section). Then, we transform the IF image into the same space as the transcriptomic image, allowing the two to be overlaid on each other. The user selects more points if necessary to obtain accurate registration as judged using the GUI.

### In vivo *to* ex vivo *image registration*

Registration of *in vivo* to *ex vivo* images is accomplished through a sequence of linear and non-linear transformations along the transform graph. The data that need to be registered consist of:

- **Functional imaging planes (two-photon *in vivo*):** Neuronal activity *in vivo* was acquired in 7 planes (six plus a flyback plane) spaced at 30µm apart. This was recorded with 920nm illumination wavelength and green emission filters to measure calcium-dependent GCaMP fluorescence.
- **Daily z-stacks (two-photon *in vivo*):** At the beginning of each recording session, a quick “daily z-stack” is collected to align that day’s functional imaging planes to the reference z-stack. We used 920 nm illumination with green emission filters to measure GCaMP fluorescence.
- **Reference z-stack (two-photon *in vivo*):** The reference z-stack is a long exposure z-stack obtained on the first day of *in vivo* imaging that defines a global coordinate system for the *in vivo* recordings. The stack is imaged twice - once at 920 nm excitation wavelength to excite GCaMP and once at 1060 nm to excite dTomato - in non-simultaneous acquisitions that must be registered to each other. Green and red emissions filter images are collected for both illumination wavelengths.
- **Fluorescent protein round (confocal *ex vivo*):** Before performing the sequential confocal imaging required for coppaFISH 3D, we stained the tissue with DAPI and performed a round of confocal imaging to measure GCaMP, dTomato, and DAPI *ex vivo* on the same microscope that will be used for coppaFISH 3D. This acquisition contains images of the same GCaMP and dTomato distribution imaged *in vivo*, which allows registration to the *in vivo* reference stack; it also contains DAPI which allows registration to *ex vivo* coppaFISH 3D sequencing, imaged with the same voxel size.
- **The anchor round of sequential imaging (confocal *ex vivo*):** At the end of sequential imaging for gene detection, we acquire an “anchor image” in which all RCPs are fluorescently labelled, which serves to define the main coordinate system for coppaFISH 3D. This image also contains DAPI, which is used to register them to the fluorescent protein round.

We register all images together in 6 steps (Supplementary Figure 6). CASTalign gives the user the option to choose a transformation for each step; in most cases, a single recommended transform worked, but for some individual experiments we needed to use different CASTalign options (for example due to strong warping); these deviations are also described. After these steps, we obtain a map from the *in vivo* functional imaging planes to their corresponding gene reads. The reference z-stack serves as a reference coordinate system, and this procedure does not require segmentation of neurons in the reference z-stack, which we found to be unreliable. The full operator and computer time taken for the registration pipeline was less than a day per mouse, including visual proofreading of all steps.

#### Step 1: Imaging plane to daily z-stack

We align each functional two-photon imaging plane to the daily z-stack, obtained in the same imaging session, using the translation and shear components of an affine transform. The GUI is initialised using a starting transform estimated for each plane through function optimisation, obtained by using differential evolution to maximise a phase-correlation objective function. The function assumed the planes were translated in 3D with respect to the daily z-stack, and also included a shear along the y-z plane, corresponding to the galvo scanning direction. These starting registrations were proofread in the CASTalign GUI. Most cases (>90%) did not need manual adjustment, but planes deep in the imaging stack with low SNR sometimes did. For these, we manually set the shear magnitude and z depth to those expected based on the mirror position readings from the microscope, and tweaked parameters in the CASTalign GUI until a match was visually evident. This procedure took less than 10 minutes per session to register all planes, including manual adjustments.

#### Step 2: Daily z-stack to reference z-stack

To register daily z-stacks to the global coordinate system defined by the reference z-stack, we used rigid registration fit by selecting about 8 keypoints, usually one in each corner of the image. This was sufficient for all but one session, which showed substantial warping, for which we additionally used a laminar triangulation following the initial rigid registration. This procedure takes less than 5 minutes per session.

#### Step 3: Reference z-stack channel registration

The reference z-stack is imaged in two acquisitions - one at 920 nm and the other at 1060 nm - which must be registered to each other. Both acquisitions include a green and red filtered image channel, which do not need registration since they are acquired simultaneously. We use the image of dTomato from the red channel of the 920 nm acquisition to register to the image of dTomato from the red channel of the 1060 nm acquisition. These are registered using a translation, determined automatically using phase correlation, which we always inspected but never needed to manually adjust. Checking the accuracy of the automated registration takes about 1 minute per mouse.

#### Step 4: Confocal channel registration

Due to our four-camera setup, all *ex vivo* confocal imaging channels must be registered to each other. This is necessary because the four cameras use the same light path, but are not perfectly aligned, showing slight rotations and translations between images. (Setups which use a single camera do not require this step.) We cannot apply the same registration learned during gene calling because a non-rigid registration is performed first in that pipeline. Confocal channel registration is performed once per section using a laminar affine transformation, and applied to both the anchor image of the sequential imaging as well as the fluorescent protein image. Alignment between cameras 1-3 is performed using the bleed-through of DAPI from the shortest wavelength laser, and on camera 4 in the anchor image by aligning DAPI to the “black holes” from the cell nuclei in the anchor image. This is performed manually once per section by selecting about 8 key points per channel, at laest one in each corner of the image, to define the laminar affine transform, and takes less than 5 minutes per section.

#### Step 5: *In vivo* reference z-stack to *ex vivo* fluorescent protein round

The main step for *in vivo* to *ex vivo* registration occurs by registering the *in vivo* reference z-stack to the fluorescent protein round, taken *ex vivo* prior to bleaching native GCaMP and dTomato. We register these images using a chain of affine transforms followed by a non-linear transform. For each step, we use the CASTalign GUI overlaying both the GCaMP and dTomato signals from both images. We normally apply coppaFISH 3D on multiple consecutive vibratome sections, which will come from neighbouring regions of 3D space *in vivo*. Furthermore, the *ex vivo* fluorescent protein image of each of these sections is composed of multiple x-y tiles. We separately register each tile of each section to the reference z-stack.

First, we manually specify an affine transform in the chain that resizes both images to have a 2µm voxel size, which provides a good trade-off between computational resources and the resolution of the section. The scaling factor is computed automatically using calibration measurements from our two-photon and confocal microscopes. Note that this resizing does not downsample or reduce the quality of the underlying image, as all transforms are simply coordinate transformations rather than image resamplings.

Next, we perform a parametric rigid transform to put the *ex vivo* tile in approximately the correct position. Because aligning the first tile of the first section is most challenging, we choose a tile whose depth and rotation can be determined using fiducial markers, typically from section near the top of the GCaMP-expressing cells, whose orientation can be found from the relative position of the dTomato stereotactic injection centroids. Aligning this tile usually takes around 30 minutes, depending on user experience. Subsequent tiles and sections can be aligned using previous tiles as a starting position; this makes registration much easier, because we know that there is a moderate overlap between the edges of tiles in the same section, and that the tiles from adjacent sections are 50µm above or below each other. Other than the first tile per mouse, as described previously, finding the rigid transformation takes about one minute per tile. This rigid registration is meant to be a rough approximation of the correct tile location.

We refine the linear fit by performing a laminar affine transformation. Since CASTalign finds transforms in real time, this is an interactive process: we iteratively add points and refit the transformation until we achieve a good fit across the tile. This generally provides a close approximation of the *in vivo* to *ex vivo* registration, but does not account for tissue warping. This step takes about 10 minutes per tile.

Finally, we account for nonlinear tissue warping using a laminar triangulation transform for each tile. This starts using the same matched pairs that defined the tile’s laminar affine transform, but the user can make finer adjustments by adding more keypoint pairs interactively. Again, because the transform is fast, this is an interactive process: the updated transformed image is redisplayed in real-time each time a keypoint pair is added. The time needed to perform this step and the refined linear fit depends greatly on the tile: some tiles do not need any nonlinear adjustments at all (in which case this step is skipped), while some require up to 30 minutes or more of extra adjustments. About 5 minutes per tile is typical.

#### Step 6: *Ex vivo* fluorescent protein image to coppaFISH 3D anchor round

Finally, we register the *ex vivo* fluorescent protein image to the coppaFISH 3D anchor round (also *ex vivo*) using an affine transform constrained to have no shear, found by point-cloud registration of nuclei detected in the DAPI channel of each image. To detect nuclei, we bandpass filter both DAPI channels with a difference of gaussians filter (0.025µm - 0.5µm), threshold at the 0.999 quantile, and construct a point cloud from the centroids of the connected components of the binary image. We perform point-cloud registration using differential evolution to globally search the space of nine transform parameters (three translations, three rotations, and three scales) for minimal squared hinge loss of the distance of each point with its nearest neighbour, normalised by the total number of points within the convex hull of both sections. We proofread this fit using the CASTalign GUI, and in the cases where the procedure fails (about 10-20% of the time, often when tissue was unmounted and remounted between the fluorescent protein image and sequential imaging) we manually align using either a laminar affine transform or a laminar triangulation. Proofreading takes about 2 minutes per tile, and fixing a tile’s registration takes about 5-10 minutes. This step completes the registration of the sequential imaging rounds of the *ex vivo* section to the imaging planes from *in vivo* two-photon imaging.

### *In vivo* to *ex vivo* cell pairing

The transform graph maps coordinates from each *in vivo* imaging plane into the coordinate system of the *ex vivo* coppaFISH 3D gene reads. To identify transcriptomically typed neurons recorded *in vivo*, we perform cell segmentation on the GCaMP channel of the *ex vivo* fluorescent protein image, and paired them with the 2D ROI suite2p cell produced by suite2p from each imaging plane. Pairing was performed in five steps:

1. **Identify repeated ROIs across recording planes**. Because we used multiplane *in vivo* calcium imaging, the same cell could appear in two or more neighbouring imaging planes. To identify duplicate ROIs, we compared all ROIs using the pairwise correlation of their calcium timecourses and the distances between their centroids. For each ROI in a non-flyback plane, candidate matches were the ROIs with the highest temporal correlation in each immediately adjacent plane; each ROI in the flyback plane was assigned a single candidate match across all other planes, because adjacency is not well defined for the flyback plane. Candidate matches were retained only if the centroid distance was less than 10 pixels (18µm) and the correlation exceeded the 95th percentile for both ROIs; duplicate ROIs were then defined as mutual best matches among these candidates. This yielded up to 7, but usually 1-3, ROIs judged to come from a single underlying cell.
2. **Quality control to exclude cells with ambiguous identity**. We excluded any cells which had ambiguous transcriptomic identity based on two criteria. First, we excluded cells which had fewer than 40 total gene reads. Second, we excluded cells which had ambiguous class. Our probabilistic cell calling algorithm gives a probability of cell identity. If the identity has at least a 10% total probability of belonging to a different class than the assigned class, or if the second highest probability cell type belonged to a different class, we excluded the cell.
3. **Transform segmented *ex vivo* cells to the imaging plane**. We matched *ex vivo* cells to *in vivo* ROIs one functional imaging plane at a time. Each imaging plane forms a two-dimensional slice through the reference z-stack, and only some *ex vivo* coppaFISH 3D tiles intersect this slice. For tiles that did, we created a silhouette image of the cells which intersected the slice, showing where each *ex vivo* GCaMP-segmented cell intersects the 2D coordinate system of imaging plane. To do this, we first dilated the 3D silhouettes of the *ex vivo* segmented cells by 6µm in the z dimension to account for the broad axial point-spread function, and then transformed them into imaging plane coordinates using the transform graph, keeping only the slice with z=0. This produced two 2D maps in the same coordinate system: one showing the ROIs from calcium imaging, and one showing the segmented *ex vivo* cells.
4. **Pairing *ex vivo* cells to ROIs**. Pairing was performed in two steps: a “pairing quality” score was computed for each candidate pair of an *in vivo* ROI and an *ex vivo* cell, and a greedy algorithm was then used to assign final pairs. To compute the score, we measured the overlap between segmented cells in the two maps. For candidates with any overlap, the score was the harmonic mean of two fractions: (a) the fraction of pixels from the *in vivo* ROI that overlapped the *ex vivo* segmented cell, and (b) the fraction of pixels from the *ex vivo* segmented cell in the plane of the *in vivo* ROI that overlapped the *in vivo* ROI. Candidate pairs were discarded if first fraction was below 50%, or the second was below 40%. From the remaining candidates, we selected the pair with the highest score and assigned it as a match. We then eliminated all other candidate pairs involving either the matched *in vivo* ROI or the matched *ex vivo* cell, and also eliminated candidate pairs involving other *in vivo* ROIs previously identified as repeated-plane views of the same cell. Iterating this procedure gave a unique mapping between segmented cells on imaging tiles and *in vivo* ROIs.
5. **Removing duplicates across neighbouring tiles**. Because adjacent tiles overlapped by 10%, an *in vivo* ROI could match *ex vivo* cells in multiple tiles if the cell was in the overlapping region. To remove these duplicates, we first checked whether the two instances of the same cell were assigned to the same transcriptomic class, since cell calling was performed independently in each tile. If so, we kept the instance with the most gene reads. If only one instance was assigned a transcriptomic identity, we used that identity. If the two instances were assigned to different classes we excluded the cell from analysis.

Following this procedure, we obtained a mapping from *in vivo* ROIs to *ex vivo* segmented cells with transcriptomic identities. Not all ROIs could be mapped to *ex vivo* cells, but each mapped ROI had a unique *ex vivo* partner. Likewise, not all *ex vivo* cells could be mapped to ROIs, but those that were mapped were assigned to a ROI, or to a set of ROIs recorded on different planes and identified as the same cell. In practice, manual inspection indicated that this procedure produced robust matches.

**Supplementary Figure 1.**
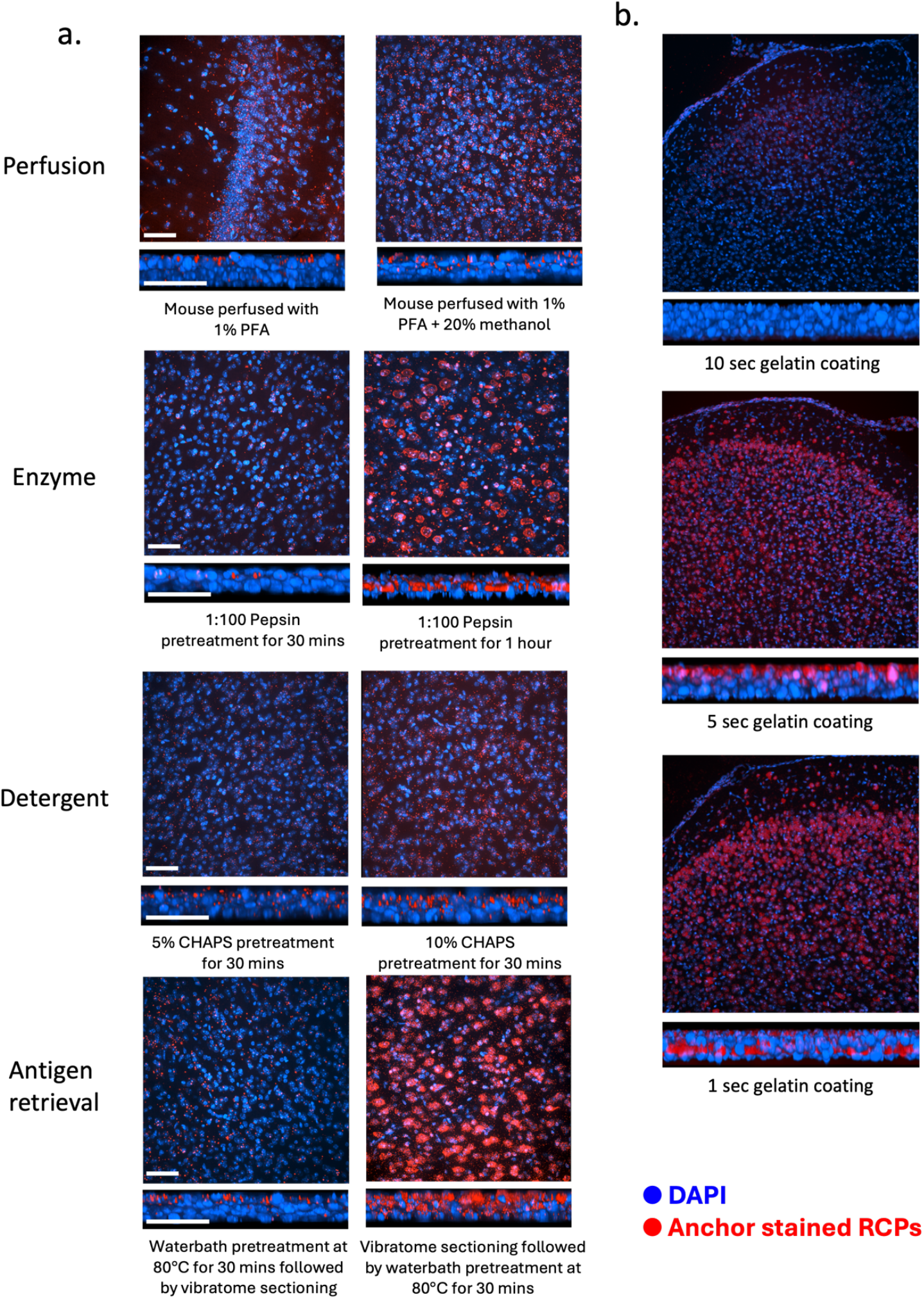
Optimisation of tissue processing conditions for coppaFISH 3D. (a) Comparison of fixation and pretreatment conditions, shown as xy images with corresponding xz views below, using DAPI (blue) and anchor-stained rolling-circle products (RCPs; red). Rows compare perfusion conditions (1% PFA versus 1% PFA + 20% methanol), pepsin pretreatment (1:100 for 30 min versus 1 h), CHAPS pretreatment (5% for 30 min versus 10% for 30 min), and heat-mediated antigen retrieval (80°C waterbath for 30 min before versus after vibratome sectioning). (b) Comparison of gelatin-coating durations used for section mounting (10 s, 5 s, and 1 s), shown with xy images and corresponding xz views, to assess tissue adherence and preservation of anchor-detected RCP signal. Blue, DAPI; red, anchor-stained RCPs.

**Supplementary Figure 2.**
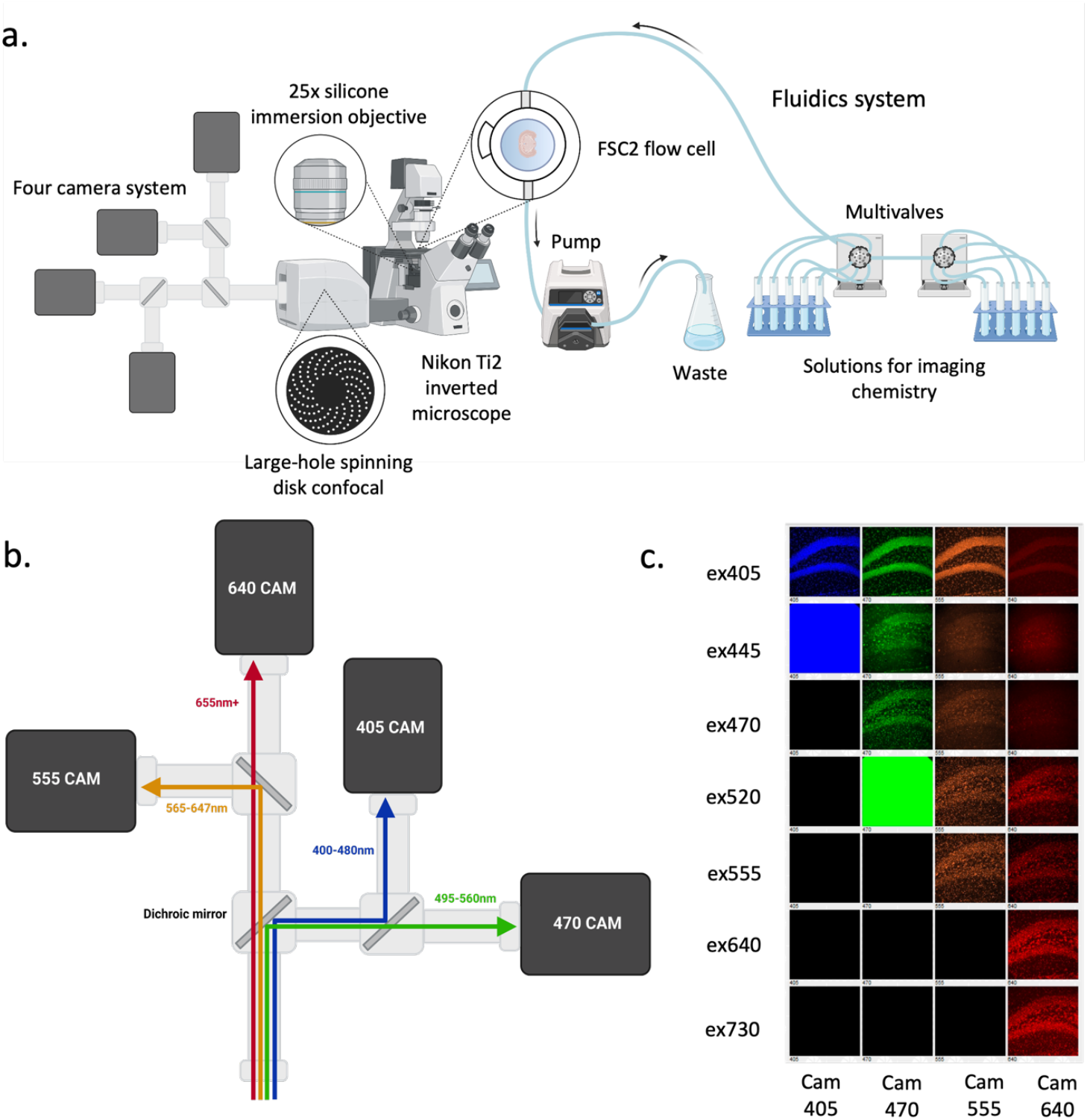
Microscope and fluidics setup for coppaFISH 3D imaging. (a) Schematic of the imaging setup, consisting of a Nikon Ti2 microscope with a large-hole spinning-disk confocal, a 25x silicone-immersion objective, a four-camera imaging system, and an FSC2 flow cell connected to an automated fluidics system for delivery of imaging chemistries. (b) Optical layout of the four-camera detection path showing spectral splitting of emitted fluorescence into the 405, 470, 555, and 640 camera channels. (c) Representative images acquired under each excitation wavelength across the four camera channels, illustrating the spectral separation used for multichannel detection. Figure created with BioRender.com.

**Supplementary Figure 3.**
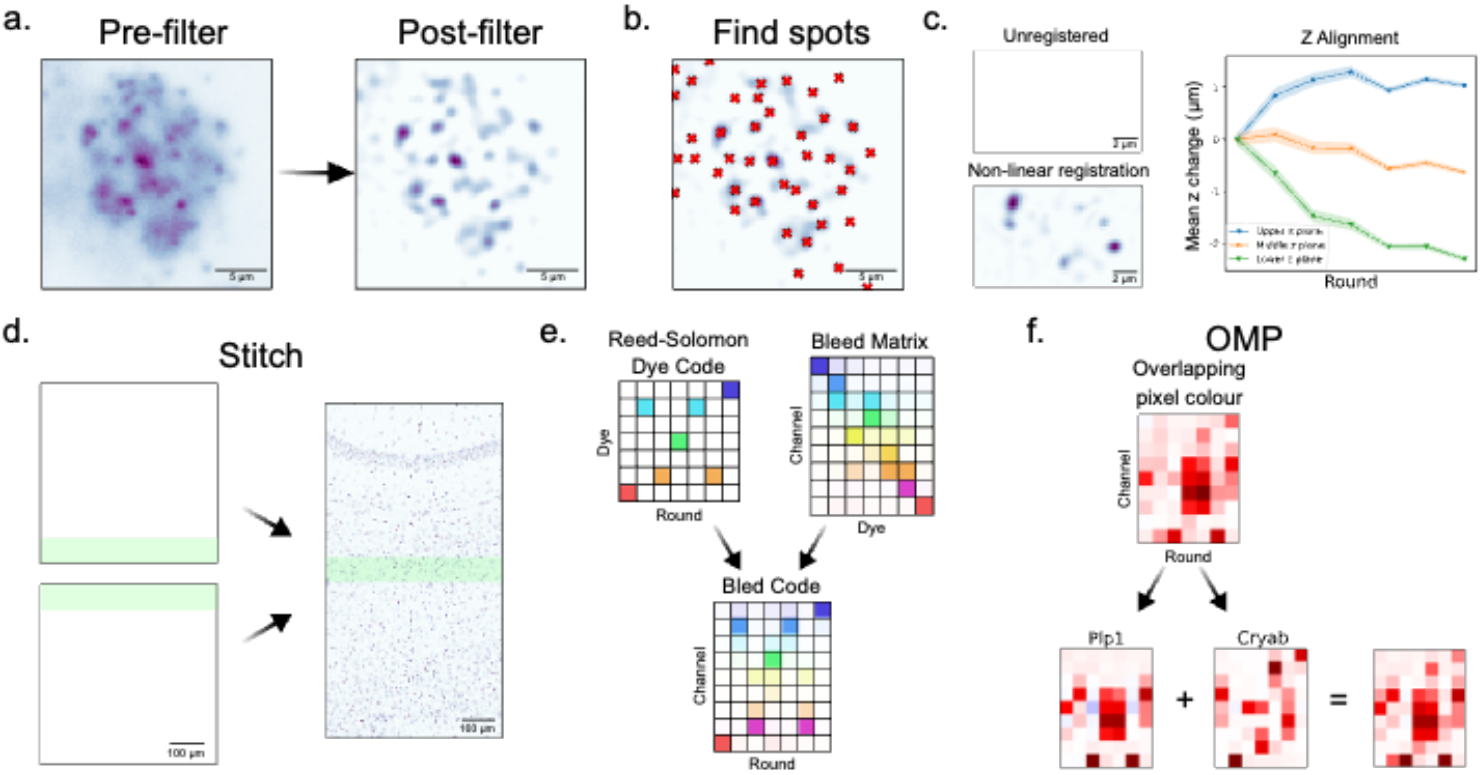
Overview of the coppafisher gene-calling pipeline. (a) Example of filtering and spot detection, showing an image before filtering, after filtering. (b) The resulting detected spot locations after filtering. (c) Example of registration quality assessment. The first column shows the voxel-wise product image for the first round before and after registration and the estimated optical flow mean z-shifts relative to the first round for upper, middle, and lower planes. (d) Stitching of adjacent fields of view into a single large image. (e) Construction of gene-specific bled codes by combining the Reed-Solomon code with the dataset-derived dye bleed matrix; example shown for Vim. (f) Illustration of OMP resolving overlapping signal, in which an observed pixel colour is decoded as a combination of two gene bled codes (example shown for Plp1 and Cryab).

**Supplementary Figure 4.**
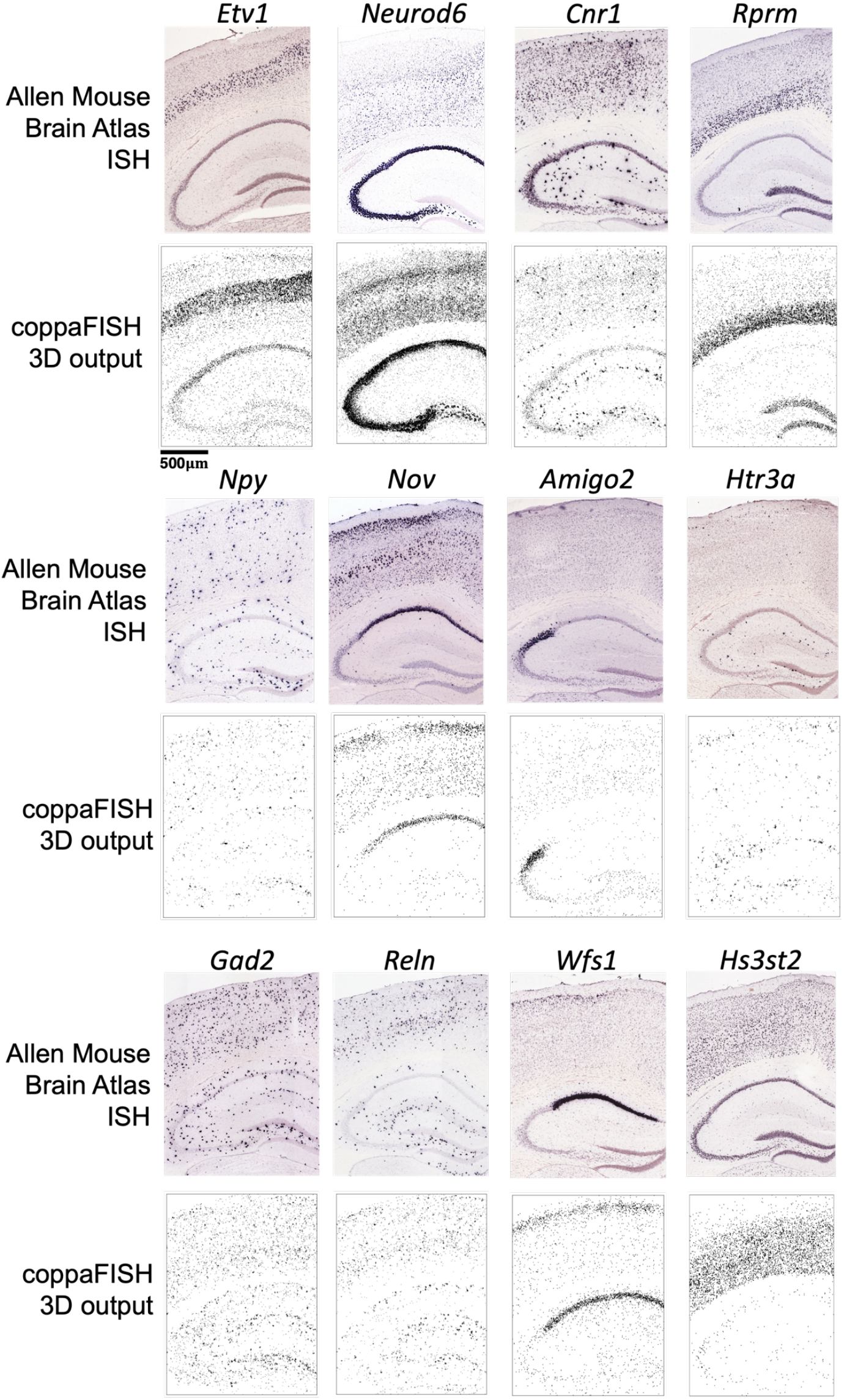
Additional comparisons between coppaFISH 3D and the Allen Mouse Brain Atlas *in situ* hybridisation reference data^41^. Spatial expression patterns for representative genes are shown in matched hippocampal and cortical regions, extending Figure 4b and illustrating the concordance between coppaFISH 3D transcript detections and published ISH atlases.

**Supplementary Figure 5.**
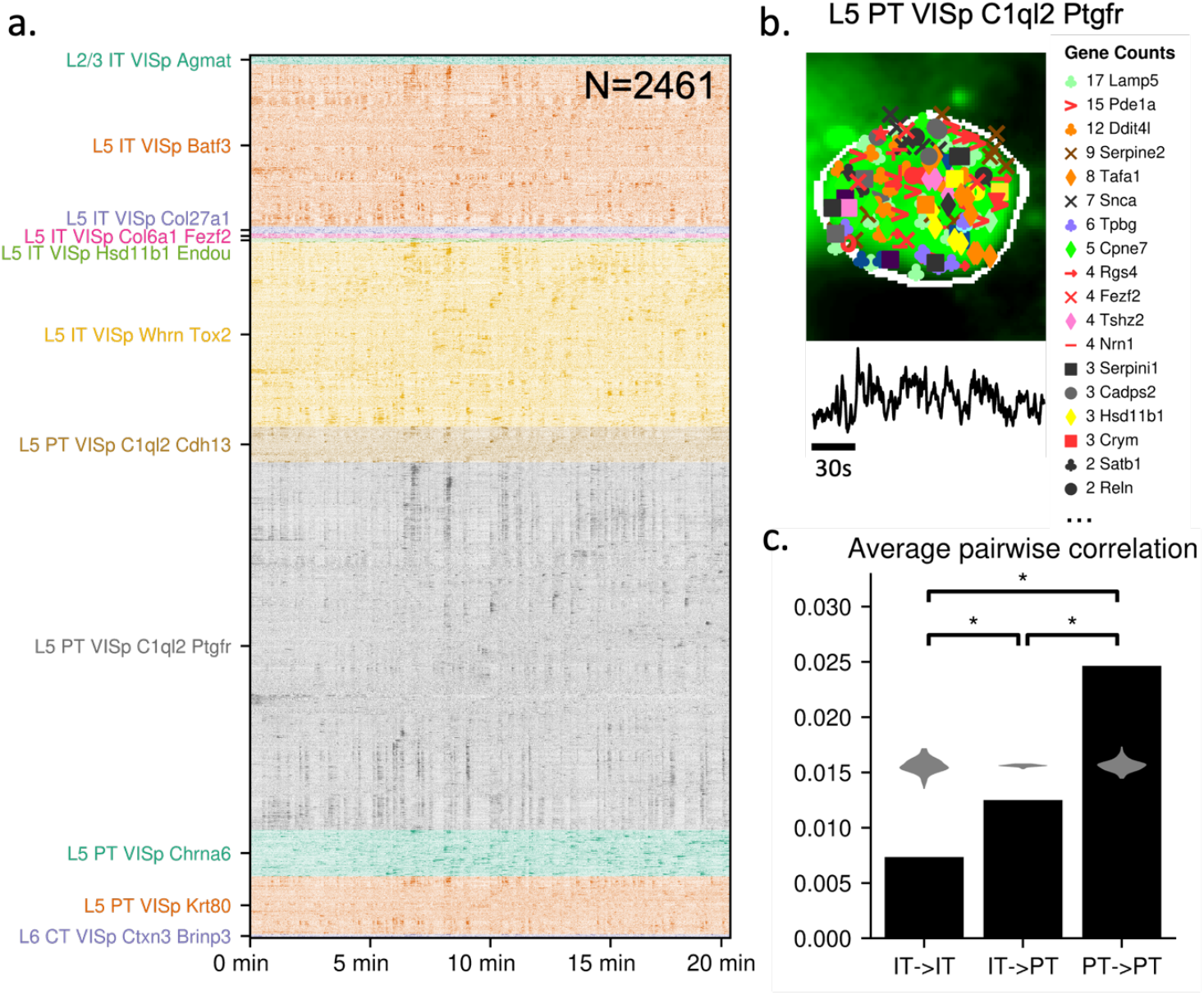
Replication of the Figure 6 activity analysis. The figure reproduces the key analyses from Figure 6d-f in an independent animal (mouse H), showing population calcium activity ordered by transcriptomic class, an example registered neuron with its assigned genes and activity trace, and pairwise-correlation comparisons among IT and PT neurons.

**Supplementary Figure 6.**
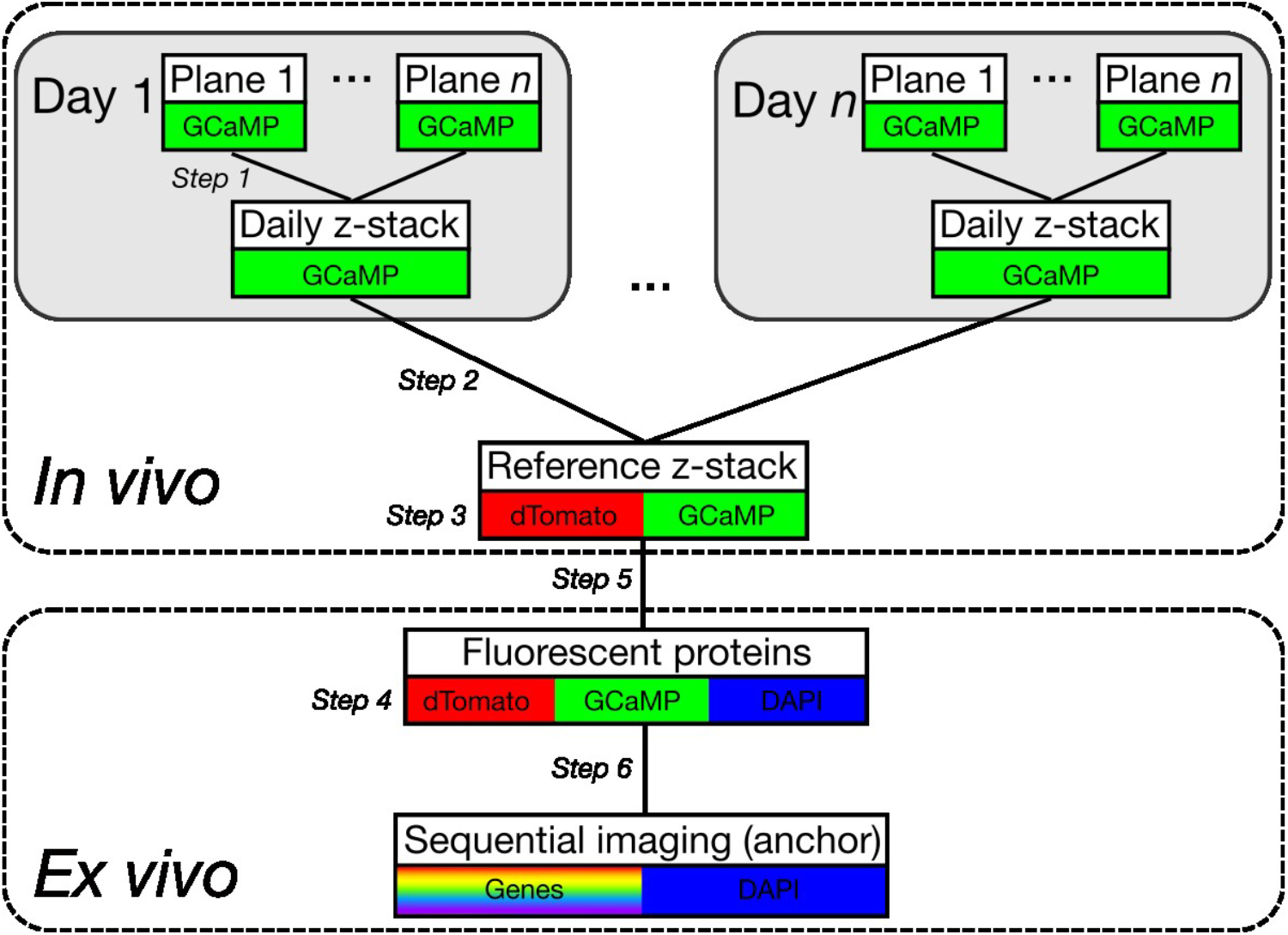
*In vivo* to *ex vivo* registration. A diagram showing the registration of imaging planes to the daily z-stack, to the reference z-stack, to the fluorescent protein image, to the anchor, to the sequential imaging. Background colours indicate the channels acquired for each image. Imaging plane ROIs are directly matched to the sequential imaging through a single transform.

**Supplementary Table 1:**
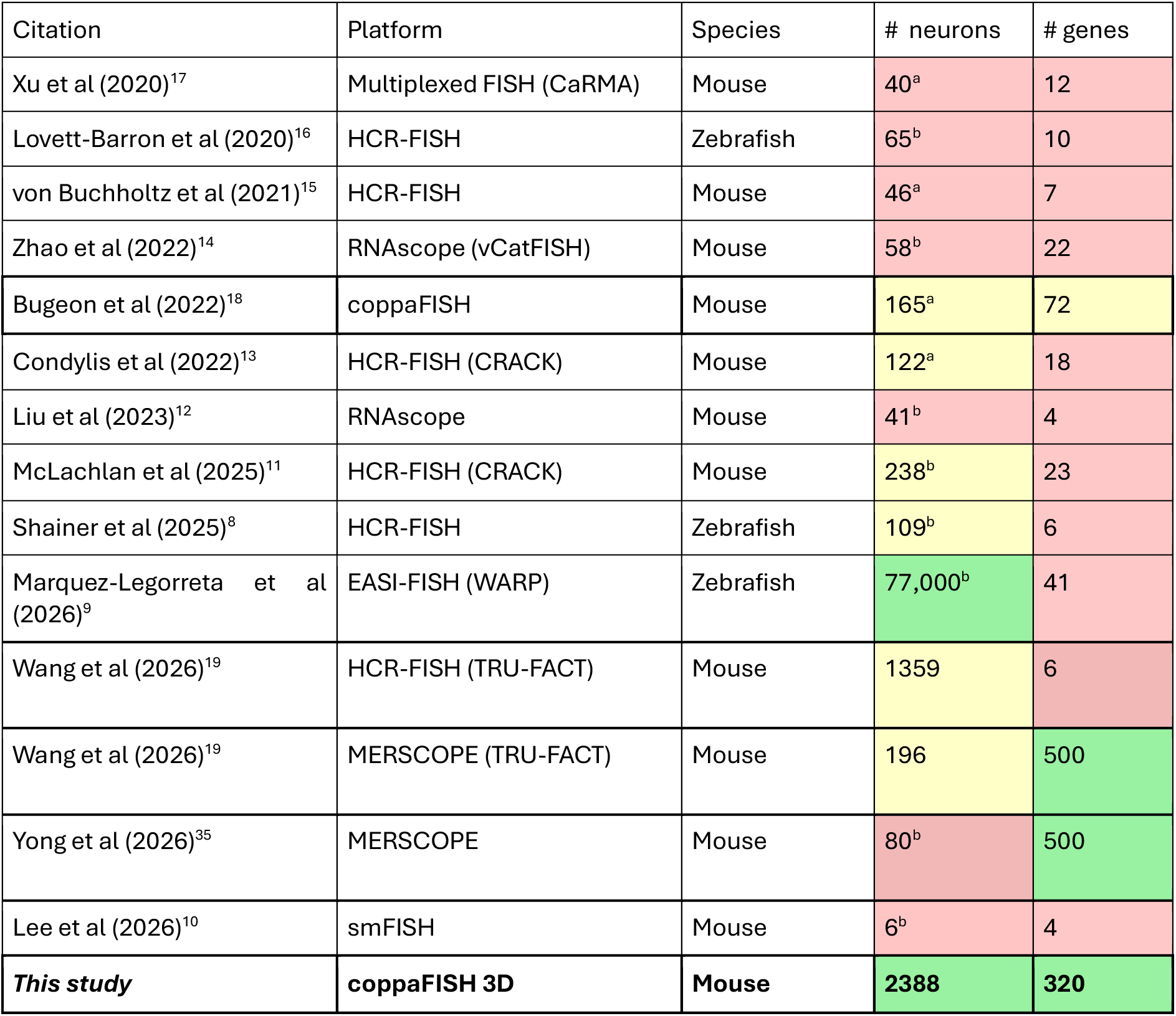
Comparison of studies pairing *in vivo* recordings with spatially resolved transcriptomics. Maximum number of simultaneously recorded neurons (“# neurons”) and maximum number of genes that were identified (“# genes”) for each study. Superscript “a” indicates the maximum reported for any session in the paper, and “b” indicates the mean reported across sessions (where individual session counts were not reported). Green indicates >1500 neurons or >100 genes, yellow indicates >100 neurons or >50 genes, and red indicates fewer than 100 neurons or 50 genes.

## Notes

### Competing Interest Statement

The authors have declared no competing interest.

### Summary of Updates

Supplemental table 1 colours fixed, grey cells were changed to green.

## References

1. Tasic, B. et al. Shared and distinct transcriptomic cell types across neocortical areas. Nature 2018 563:7729 563, 72–78 (2018).

2. Scala, F. et al. Phenotypic variation of transcriptomic cell types in mouse motor cortex. Nature 2020 598:7879 598, 144–150 (2020).

3. Gouwens, N. W. et al. Classification of electrophysiological and morphological neuron types in the mouse visual cortex. Nature Neuroscience 2019 22:7 22, 1182–1195 (2019).

4. Kalmbach, B. E. et al. Signature morpho-electric, transcriptomic, and dendritic properties of human layer 5 neocortical pyramidal neurons. Neuron 109, 2914–2927.e5 (2021).

5. Bugeon, S. et al. A transcriptomic axis predicts state modulation of cortical interneurons. Nature 2022 607:7918 607, 330–338 (2022).

6. Zhang, M. et al. Spatially resolved cell atlas of the mouse primary motor cortex by MERFISH. Nature 2021 598:7879 598, 137–143 (2021).

7. Green, J. et al. A cell-type-specific error-correction signal in the posterior parietal cortex. Nature 2023 620:7973 620, 366–373 (2023).

8. Shainer, I. et al. Transcriptomic neuron types vary topographically in function and morphology. Nature 638, 1023–1033 (2025).

9. Marquez-Legorreta, E. et al. Whole-Brain Co-Mapping of Gene Expression and Neuronal Activity at Cellular Resolution in Behaving Zebrafish. bioRxiv doi:10.64898/2026.02.07.704095. 2026.02.07.704095 (2026)

10. Lee, S. et al. Correlative Imaging Platform Linking Taste Cell Function to Molecular Identity. Advanced Science 13, e11309 (2026).

11. McLachlan, C. A. et al. Transcriptional determinants of goal-directed learning and representational drift in the parahippocampal cortex. Cell Rep. 44, (2025).

12. Liu, Y. et al. Mapping visual functions onto molecular cell types in the mouse superior colliculus. Neuron 111, 1876–1886.e5 (2023).

13. Condylis, C. et al. Dense functional and molecular readout of a circuit hub in sensory cortex. Science 375, (2022).

14. Zhao, Q. et al. A multidimensional coding architecture of the vagal interoceptive system. Nature 603, 878–884 (2022).

15. von Buchholtz, L. J. et al. Decoding Cellular Mechanisms for Mechanosensory Discrimination. Neuron 109, 285–298.e5 (2021).

16. Lovett-Barron, M. et al. Multiple convergent hypothalamus-brainstem circuits drive defensive behavior. Nat. Neurosci. 23, 959–967 (2020).

17. Xu, S. et al. Behavioral state coding by molecularly defined para-ventricular hypothalamic cell type ensembles. Science 370, (2020).

18. Bugeon, S. et al. A transcriptomic axis predicts state modulation of cortical interneurons. Nature 607, 330–338 (2022).

19. Wang, L. et al. Multimodal alignments of in vivo imaging and spatial biology datasets at cellular resolution. bioRxiv 2026.04.28.719500 (2026) doi:10.64898/2026.04.28.719500.

20. Kalhor, K. et al. Mapping human tissues with highly multiplexed RNA in situ hybridization. Nature Communications 2024 15:1 15, 2511– (2024).

21. Lee, J. H. et al. Highly multiplexed subcellular RNA sequencing in situ. Science (1979). 343, 1360–1363 (2014).

22. Rouhanifard, S. H. et al. ClampFISH detects individual nucleic acid molecules using click chemistry–based amplification. Nature Biotechnology 2018 37:1 37, 84–89 (2018).

23. Wang, Y. et al. EASI-FISH for thick tissue defines lateral hypothalamus spatio-molecular organization. Cell 184, 6361–6377.e24 (2021).

24. Codeluppi, S. et al. Spatial organization of the somatosensory cortex revealed by osmFISH. Nature Methods 2018 15:11 15, 932–935 (2018).

25. Lubeck, E. & Cai, L. Single-cell systems biology by super-resolution imaging and combinatorial labeling. Nature Methods 2012 9:7 9, 743–748 (2012).

26. Zeng, H. et al. Integrative in situ mapping of single-cell transcriptional states and tissue histopathology in a mouse model of Alzheimer’s disease. Nature Neuroscience 2023 26:3 26, 430–446 (2023).

27. Wang, X. et al. Three-dimensional intact-tissue sequencing of single-cell transcriptional states. Science (1979). 361, (2018).

28. Gyllborg, D. et al. Hybridization-based in situ sequencing (HybISS) for spatially resolved transcriptomics in human and mouse brain tissue. Nucleic Acids Res. 48, e112–e112 (2020).

29. Ke, R. et al. In situ sequencing for RNA analysis in preserved tissue and cells. Nature Methods 2013 10:9 10, 857–860 (2013).

30. Choi, H. M. T. et al. Programmable in situ amplification for multiplexed imaging of mRNA expression. Nature Biotechnology 2010 28:11 28, 1208–1212 (2010).

31. Borm, L. E. et al. Scalable in situ single-cell profiling by electrophoretic capture of mRNA using EEL FISH. Nature Biotechnology 2022 41:2 41, 222–231 (2022).

32. Moffitt, J. R. et al. High-throughput single-cell gene-expression profiling with multiplexed error-robust fluorescence in situ hybridization. Proc. Natl. Acad. Sci. U. S. A. 113, 11046–11051 (2016).

33. Wang, F. et al. RNAscope: A Novel in Situ RNA Analysis Platform for Formalin-Fixed, Paraffin-Embedded Tissues. The Journal of Molecular Diagnostics 14, 22–29 (2012).

34. Femino, A. M., Fay, F. S., Fogarty, K. & Singer, R. H. Visualization of single RNA transcripts in situ. Science (1979). 280, 585–590 (1998).

35. Choong Yong, H. et al. A transcriptomic axis aligns with in vivo functional dynamics in hippocampal inhibitory circuits. bioRxiv 2026.04.07.716935 (2026) doi:10.64898/2026.04.07.716935.

36. Stringer, C., Wang, T., Michaelos, M. & Pachitariu, M. Cellpose: a generalist algorithm for cellular segmentation. Nature Methods 2020 18:1 18, 100–106 (2020).

37. Qian, X. et al. Probabilistic cell typing enables fine mapping of closely related cell types in situ. Nat. Methods 17, 101–106 (2020).

38. Yao, Z. et al. A high-resolution transcriptomic and spatial atlas of cell types in the whole mouse brain. Nature 2023 624:7991 624, 317–332 (2023).

39. Tasic, B. et al. Shared and distinct transcriptomic cell types across neocortical areas. Nature 2018 563:7729 563, 72–78 (2018).

40. ISH Data:: Allen Brain Atlas: Mouse Brain. http://mouse.brain-map.org/search/index.

41. Foiani, M. S. et al. Distinct mechanistic pathways of early tauopathy revealed by MAPT mutations. bioRxiv 2026.02.14.705716 (2026) doi:10.64898/2026.02.14.705716.

42. Hsu, J., Nguyen, K. T., Bujnowska, M., Janes, K. A. & Fallahi-Sichani, M. Protocol for iterative indirect immunofluorescence imaging in cultured cells, tissue sections, and metaphase chromosome spreads. STAR Protoc. 5, 103190 (2024).

43. Mallach, A. et al. Microglia-astrocyte crosstalk in the amyloid plaque niche of an Alzheimer’s disease mouse model, as revealed by spatial transcriptomics. Cell Rep. 43, (2024).

44. Chen, W. T. et al. Spatial Transcriptomics and In Situ Sequencing to Study Alzheimer’s Disease. Cell 182, 976–991.e19 (2020).

45. Fang, R. et al. Three-dimensional single-cell transcriptome imaging of thick tissues. Elife 12, (2024).

46. Schulte, S. J., Fornace, M. E., Hall, J. K., Shin, G. J. & Pierce, N. A. HCR spectral imaging: 10-plex, quantitative, high-resolution RNA and protein imaging in highly autofluorescent samples. Development (Cambridge) 151, (2024).

47. Asami, S., Yin, C., Garza, L. A. & Kalhor, R. Deconvolving organogenesis in space and time via spatial transcriptomics in thick tissues. bioRxiv 2024.09.24.614640 (2024) doi:10.1101/2024.09.24.614640.

48. Blackman, R. B. & Tukey, J. W. The Measurement of Power Spectra from the Point of View of Communications Engineering. (New York: Dover Publications, 1959).

49. Wiener, N. Extrapolation, Interpolation, and Smoothing of Stationary Time Series: With Engineering Applications. Extrapolation, Interpolation, and Smoothing of Stationary Time Series https://doi.org/10.7551/MITPRESS/2946.001.0001 (1949) doi:10.7551/MITPRESS/2946.001.0001.

50. Lucas, B. D. & Kanade, T. An Iterative Image Registration Technique with an Application to Stereo Vision.

51. Arun, K. S., Huang, T. S. & Blostein, S. D. Least-Squares Fitting of Two 3-D Point Sets. IEEE Trans. Pattern Anal. Mach. Intell. PAMI-9, 698–700 (1987).

52. Thurman, S. T., Guizar-Sicairos, M. & Fienup, J. R. Efficient subpixel image registration algorithms. Optics Letters, Vol. 33, Issue 2, pp. 156-158 33, 156–158 (2008).

53. Skriabine, S., Shinn, M., Picard, S., Harris, K. D. & Carandini, M. Mapping the visual cortex with Zebra noise and wavelets. J. Vis. 26, (2026).

54. Bogovic, J. A., Hanslovsky, P., Wong, A. & Saalfeld, S. Robust registration of calcium images by learned contrast synthesis. Proceedings - International Symposium on Biomedical Imaging 2016-June, 1123–1126 (2016).

